# Analysis of the Diverse Antigenic Landscape of the Malaria Invasion Protein RH5 Identifies a Potent Vaccine-Induced Human Public Antibody Clonotype

**DOI:** 10.1101/2023.10.04.560576

**Authors:** Jordan R. Barrett, Dimitra Pipini, Nathan D. Wright, Andrew J. R. Cooper, Giacomo Gorini, Doris Quinkert, Amelia M. Lias, Hannah Davies, Cassandra Rigby, Maya Aleshnick, Barnabas G. Williams, William J. Bradshaw, Neil G. Paterson, Thomas Martinson, Payton Kirtley, Luc Picard, Christine D. Wiggins, Francesca R. Donnellan, Lloyd D. W. King, Lawrence T. Wang, Jonathan F. Popplewell, Sarah E. Silk, Jed de Ruiter Swain, Katherine Skinner, Vinayaka Kotraiah, Amy R. Noe, Randall S. MacGill, C. Richter King, Ashley J. Birkett, Lorraine A. Soisson, Angela M. Minassian, Douglas A. Lauffenburger, Kazutoyo Miura, Carole A. Long, Brandon K. Wilder, Lizbé Koekemoer, Joshua Tan, Carolyn M. Nielsen, Kirsty McHugh, Simon J. Draper

**Author notes:** These authors contributed equally (Kirsty McHugh, Simon J. Draper).

## Abstract

The highly conserved and essential *Plasmodium falciparum* reticulocyte-binding protein homolog 5 (PfRH5) has emerged as the leading target for vaccines that seek to protect against the disease-causing blood-stage of malaria. However, the features of the human vaccine-induced antibody response that confer highly potent inhibition of malaria parasite invasion into red blood cells are not well defined. Here we characterize over 200 human IgG monoclonal antibodies induced by the most advanced PfRH5 vaccine. We define the antigenic landscape of this molecule, and establish epitope specificity, antibody association rate and intra-PfRH5 antibody interactions are key determinants of functional anti-parasitic potency. In addition, we identify a germline gene combination that results in an exceptionally potent class of antibody and demonstrate its prophylactic potential to protect against *P. falciparum* parasite challenge *in vivo*. This comprehensive dataset provides a framework to guide rational design of next-generation vaccines and prophylactic antibodies to protect against blood-stage malaria.

## INTRODUCTION

The exponential growth of *Plasmodium falciparum* parasites in the blood of infected individuals drives the disease state known as malaria. Preventative measures including the use of insecticides, bednets and anti-malarial drugs have proven effective in reducing the global malaria burden since the turn of the millennium, however recent evidence suggests progress has stalled. Current estimates indicate clinical cases rose in 2021 to 247 million, leading to ∼619,000 deaths, primarily in young African infants and children ^1^. Immune-based interventions, including prophylactic monoclonal antibodies (mAbs) and vaccines, offer highly promising and alternative strategies to complement the current public health tools for malaria ^2,3^. Indeed, substantial progress has been made recently with the clinical development of the RTS,S/AS01 and R21/Matrix-M™ subunit vaccines and the L9LS mAb, all of which target the *P. falciparum* circumsporozoite protein (PfCSP) thereby blocking infection at the pre-erythrocytic stage ^4–6^. Nevertheless, when these interventions fail or immunity wanes, sporozoites eventually establish liver infection from where they develop into merozoites that emerge to initiate their continual and disease-causing cycle of growth in the blood. Blockade of merozoite invasion into the host red blood cell (RBC) thus provides a second and highly complementary opportunity for immune-intervention. Vaccines or mAbs against blood-stage antigens could provide standalone immunity, but also offer a leading strategy to achieve very high and more durable efficacy against *P. falciparum* via combination with the anti-PfCSP interventions in a multi-stage approach.

Merozoite invasion of the human RBC is a rapid and complex process, mediated by numerous host receptor – parasite ligand interactions. Indeed, the polymorphic and redundant nature of these parasitic targets thwarted blood-stage vaccine development for many years ^2^, however, the discovery of the *P. falciparum* reticulocyte-binding protein homolog 5 (PfRH5) as a target that overcame these historical challenges has offered new promise ^7,8^. PfRH5 is highly conserved and presented on the parasite’s apical surface within a pentameric invasion complex ^9,10^; here it forms an essential interaction with host basigin on the RBC ^8^ that also defines the human host tropism of this parasite ^11^. Vaccination studies with PfRH5 in *Aotus* monkeys and healthy UK adults have shown significant efficacy against blood-stage challenge with *P. falciparum*, with the degree of *in vivo* inhibition of parasite growth strongly correlating with *in vitro* growth inhibition activity (GIA) as measured using purified total IgG in a standardized assay ^12,13^. This ability of vaccine-induced anti-PfRH5 growth inhibitory antibodies to protect against blood-stage *P. falciparum* was subsequently validated by passive transfer of mAb in both *Aotus* monkeys ^14^ and humanized mice ^15^. Nonetheless, the protection outcomes in the UK adult clinical trial, of a protein-in-adjuvant vaccine called RH5.1/AS01_B_, were relatively modest ^13^. These data thus indicated a clear need to increase the quantitative magnitude and/or qualitative potency of the vaccine-induced anti-PfRH5 polyclonal IgG response by next-generation vaccines in order to reach the same high levels of protection observed in the *Aotus* monkey model ^12^.

The RH5.1 soluble protein vaccine ^16^, as well as another PfRH5 viral-vectored vaccine ^17,18^, are in Phase 1/2 clinical trials and deliver the full-length PfRH5 molecule (526 amino acids; ∼60 kDa in size). *In silico* analyses initially indicated regions of disorder within full-length PfRH5 including a long N-terminal region, an intrinsic loop and a small C-terminus. Thereafter, a crystal structure was first reported using a protein known as RH5ΔNL that included the small C-terminus but lacked the disordered N-terminus and intrinsic disordered loop (IDL); this showed an α-helical diamond-like architecture forms the core of the PfRH5 protein, with basigin binding across the top of the diamond-like molecule ^19^. At the bottom of the diamond, PfRH5 forms an interaction with the *P. falciparum* cysteine-rich protective antigen (PfCyRPA), thereby joining it to the hetero-pentameric invasion complex that displays PfRH5 towards the RBC membrane ^9,20^.

Further studies have since investigated individual or small panels of anti-PfRH5 mAbs raised in mice ^21–23^ or from humans vaccinated with the first-generation viral-vectored vaccine ^24–26^. These studies have provided valuable insights but lacked sufficient power to understand the relationships that underlie human antibody recognition of PfRH5 and functional growth inhibition of *P. falciparum*. We therefore conducted a high-throughput campaign to isolate over 200 novel anti-PfRH5 human mAbs from vaccinees in the RH5.1/AS01_B_ vaccine trial who showed reduced growth of blood-stage *P. falciparum* following experimental challenge ^13^. Characterization of this large panel of new clones defines the determinants of antibody functional potency across a varied epitope landscape, thereby providing the high-resolution data needed for next-generation PfRH5 vaccine immunogen design. In addition, we identify a germline gene combination that results in an exceptionally potent class of anti-PfRH5 antibody and demonstrate its prophylactic potential to protect against *P. falciparum* parasite burden *in vivo*.

## RESULTS

### The functional epitope landscape of PfRH5

Peripheral blood mononuclear cells (PBMC) were collected from UK adult volunteers vaccinated with a full-length PfRH5 soluble protein vaccine, called RH5.1, formulated in AS01_B_ adjuvant ^13,16^. PfRH5-specific IgG+ B cells were sorted by fluorescence assisted cell sorting (FACS) using a probe bound to streptavidin labelled with two different fluorophores (**Figure S1A**). This probe was composed of a biotinylated form of PfRH5 lacking the disordered N- and C-termini (called “RH5ΔNC”), given we have reported that these regions do not contribute growth-inhibitory antibodies in the vaccinees’ sera. Cells that were double-positive for both probes were sorted and lysed. Matched heavy and light chain variable antibody gene sequences were obtained through reverse transcriptase PCR (RT-PCR). Antibody genes were cloned into vectors encoding the human IgG1 backbone for expression in HEK293T cells and purification of recombinant mAbs. Purified mAbs were screened by ELISA for binding to RH5.1, resulting in the isolation of 236 novel anti-PfRH5 mAb clones. Of these, nine (3.8%) were capable of binding heat-denatured RH5.1, indicating that they bind a linear epitope (**Figure S1B**). To map these linear epitopes, the mAbs were then tested for binding to a panel of 62 previously reported 20-mer overlapping peptides spanning the length of PfRH5 ^17^ (**Figure S1C**); 7/9 mAbs bound at least one peptide in the panel. The majority of these (6/7) bound peptides 27-34, corresponding to the IDL region of PfRH5; the remaining mAb bound peptide 35 (clone R5.246). This peptide borders the IDL and is the only linear epitope mapped that also lies within the conformational core of PfRH5 (RH5ΔNL).

To further resolve the epitopes of all antibodies, the 236 mAb panel was subjected to competition binning on RH5.1 using high-throughput surface plasmon resonance (HT-SPR). These studies used seven human “sentinel mAbs” from a previous study ^24^ to bridge this work to the epitope communities identified here in this much larger mAb panel. Thirty mAbs were excluded from this analysis due to behaviour incompatible with the assay (**Data S1A**). The remaining 206 mAbs (+7 sentinels) were sorted into epitope communities using Carterra Epitope software. Data for seven mAbs required manual processing, otherwise normalized response unit (RU) values for every other mAb pair were automatically sorted into a heatmap readout (**Figure S1D, Data S1B**). These values were then used to plot a dendrogram (**Figure S1E**) to cluster antibodies, and a cut-off height was set to classify mAbs into monophyletic communities. These designations were overlaid onto a community network plot, to visualize the mAb communities and their competitive binding interactions (**Figure 1A**). This analysis resulted in the definition of 12 epitope communities: 1a (Blue, N=75, sentinel R5.004); 1b (Cyan, N=6); 1c (Grey, N=1); 2 (Red, N=43, sentinel R5.016); 3a (Violet, N=14); 3b (Pink, N=17, sentinel R5.008); 4a (Teal, N=10); 4b (Green; N=3, sentinel R5.011); 4c (Turquoise, N=2); 5a (Orange, N=25, sentinel R5.015); and 5b (Yellow, N=4, sentinel R5.001). The IDL binders were pooled together in community 6 (the group that required manual processing); these span a series of adjacent linear epitopes as defined above (Purple, N=6, sentinel R5.007). Several of these communities (identified by subletters a-c) are further grouped into supercommunities 1, 3, 4 and 5, given they share overlapping competition profiles, but differ in their competition with external communities (**Figure 1A**, **Data S1C**). Notably, the human anti-PfRH5 mAbs isolated in this study were predominantly from epitope communities 1a (Blue = 75/206) and 2 (Red = 43/206), together comprising over half of the panel. Moreover, communities 1b (Cyan), 1c (Grey), 3a (Violet), 4a (Teal) and 4c (Turquoise) represent novel epitope regions of the PfRH5 molecule that are recognized by human mAbs that were not identified by the sentinel mAbs.

**Figure 1:**
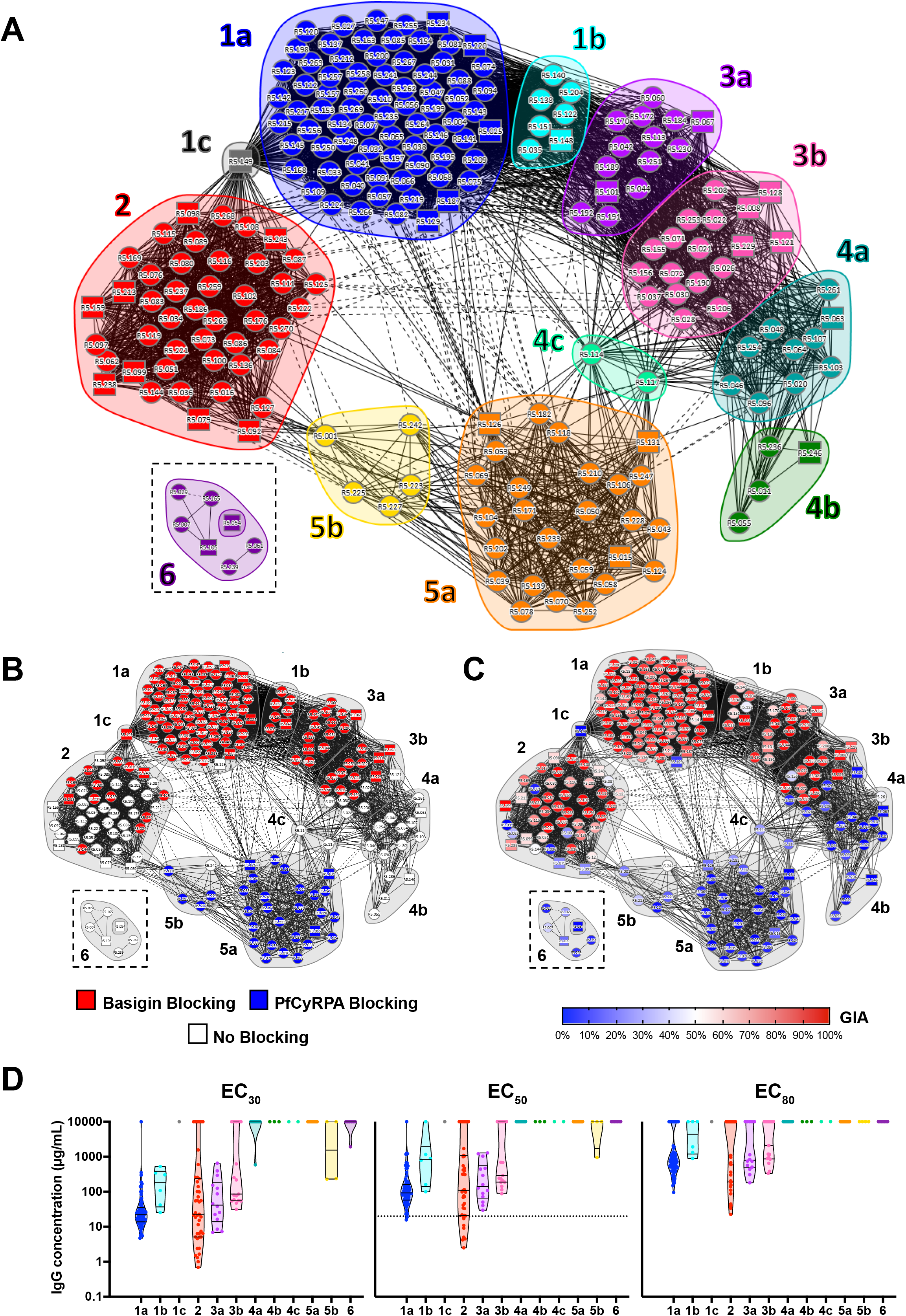
The functional epitope landscape of PfRH5. (**A**) Community network plot illustrating the competitive relationship between 206 vaccine-induced anti-PfRH5 human mAbs. Supercommunities and communities are defined by number code and color. Individual mAbs are represented as nodes. Solid lines between nodes indicate bidirectional competition. Dashed lines between nodes indicate unidirectional competition. Square nodes indicate mAbs that were excluded as either a ligand or an analyte. Community 6 (representing the IDL binders; N=7) was analysed separately and is shown here as an inset. (**B**) Community network plot overlaid with blocking category for PfRH5 binding to basigin or PfCyRPA as defined by BLI, or (**C**) GIA % as tested using a high concentration (0.8-2 mg/mL) of each mAb. (**D**) Violin plots showing the GIA potency of each epitope community as measured by the effective concentration (EC) needed to reach 30 %, 50 % or 80 % GIA. Data are log transformed and lines indicate the median and quartiles. Dashed line indicates the previously reported best-in-class sentinel mAb, R5.016 ^24^, with GIA EC_50_ against 3D7 clone *P. falciparum* = 20.7 µg/mL (**Data S1C**). Weak or non-active mAbs, for which EC values could not be determined, were assigned values of 10 mg/mL for the purpose of analysis.

To understand the functionality of these epitope communities, we next determined the ability of each mAb to block binding of PfRH5 to basigin and PfCyRPA by bio-layer interferometry (BLI). These binary blocking results were then overlaid onto the community network plot (**Figure 1B**). Antibodies in supercommunity 1 and community 3a blocked basigin binding, along with a fraction of those in communities 2 (13/43) and 3b (11/17). These latter two communities likely lie on the edge of the basigin binding site on PfRH5, explaining their bimodal functionality. PfCyRPA-blocking mAbs were entirely confined to supercommunity 5.

We next measured the ability of mAbs to block 3D7 clone *P. falciparum* parasite invasion *in vitro* (at a high concentration of 0.8-2 mg/mL) using the assay of GIA, and also overlaid these data onto the community network plot (**Figure 1C**). The distribution of GIA-positive mAbs largely followed that of the basigin-blocking mAbs, although community 2 showed a subset of growth inhibitory antibodies that did not block basigin binding as measured by BLI. Conversely, community 1c (clone R5.149) blocked basigin binding but did not show evidence of GIA. Comparison of the GIA of basigin-blocking and non-blocking clones demonstrated a mutually exclusive and significant relationship (*P*<0.0001, Mann-Whitney test) in supercommunities 1 and 3 (with only two exceptions, R5.149 and R5.030) (**Figure S1F**). In contrast, the same analysis with community 2 mAbs revealed no significant relationship between GIA and basigin-blocking as measured by BLI (**Figure S1G**). To measure relative GIA potency, mAbs were subsequently tested by GIA assay using a dilution series. Non-linear regression was fitted to the resultant log-transformed data to interpolate the effective concentrations needed to reach 30 % (EC_30_), 50 % (EC_50_) and 80 % (EC_80_) GIA (**Figure 1D, Data S1D**). The most potent mAbs were found in communities 1a, 2 and 3a, however, the spread of GIA potency within these differed. Notably, most members of community 1a had a similar potency, whilst community 2 showed a much wider distribution, ranging from multiple GIA-negative antibodies through to the most potent in the entire panel. Indeed, the most potent mAb in community 2, R5.034, showed an EC_50_ of 2.5 µg/mL, 8.3-fold lower than the previously reported best-in-class (mAb R5.016, also the sentinel for community 2) ^24^.

### Antibody on-rate correlates with growth inhibitory activity

To further investigate contributing factors to GIA potency, the binding kinetics of all 206 mAbs (+7 sentinels) to full-length RH5.1 protein were determined using HT-SPR (**Data S2**). An iso-affinity plot of mAb association (*K*_on_) and dissociation (*K*_off_) rates revealed a range of antibody affinity constants (K_D_) between 30 pM – 10 nM and an average K_D_ of approximately 1 nM (**Figure 2A**). All epitope communities spanned a similar range of affinity constants (**Figure S2A**). Fifty-eight mAbs (27%) had *K*_off_ rates too slow to reliably determine within the parameters of the assay (<6 x 10^-5^ s^-1^). R5.034 in community 2, the most potent mAb identified in the assay of GIA, had one of the fastest *K*_on_ values (1.69 x 10^6^ M^-1^ s^-1^) and had a dissociation rate constant too slow to be reported under the experimental conditions used. We also analysed the kinetic data according to the RH5.1/AS01_B_ vaccine dosing regimen in the Phase 1/2a clinical trial. These mAbs were isolated from vaccinees after their final immunization, either given in a monthly dosing schedule, or following a delayed (4-5 month) final booster dose; we previously reported the latter groups showed higher anti-RH5.1 polyclonal IgG avidity by ELISA ^13^. Notably, mAbs derived from vaccinees receiving the delayed booster doses had highly significantly different *K*_off_ and K_D_ values as compared to those receiving a monthly boost (**Figure S2B-D**), suggesting that delayed boosting in a human vaccination regimen can substantially impact the affinity of the resultant antibody response and this is largely driven by slower dissociation rates.

**Figure 2:**
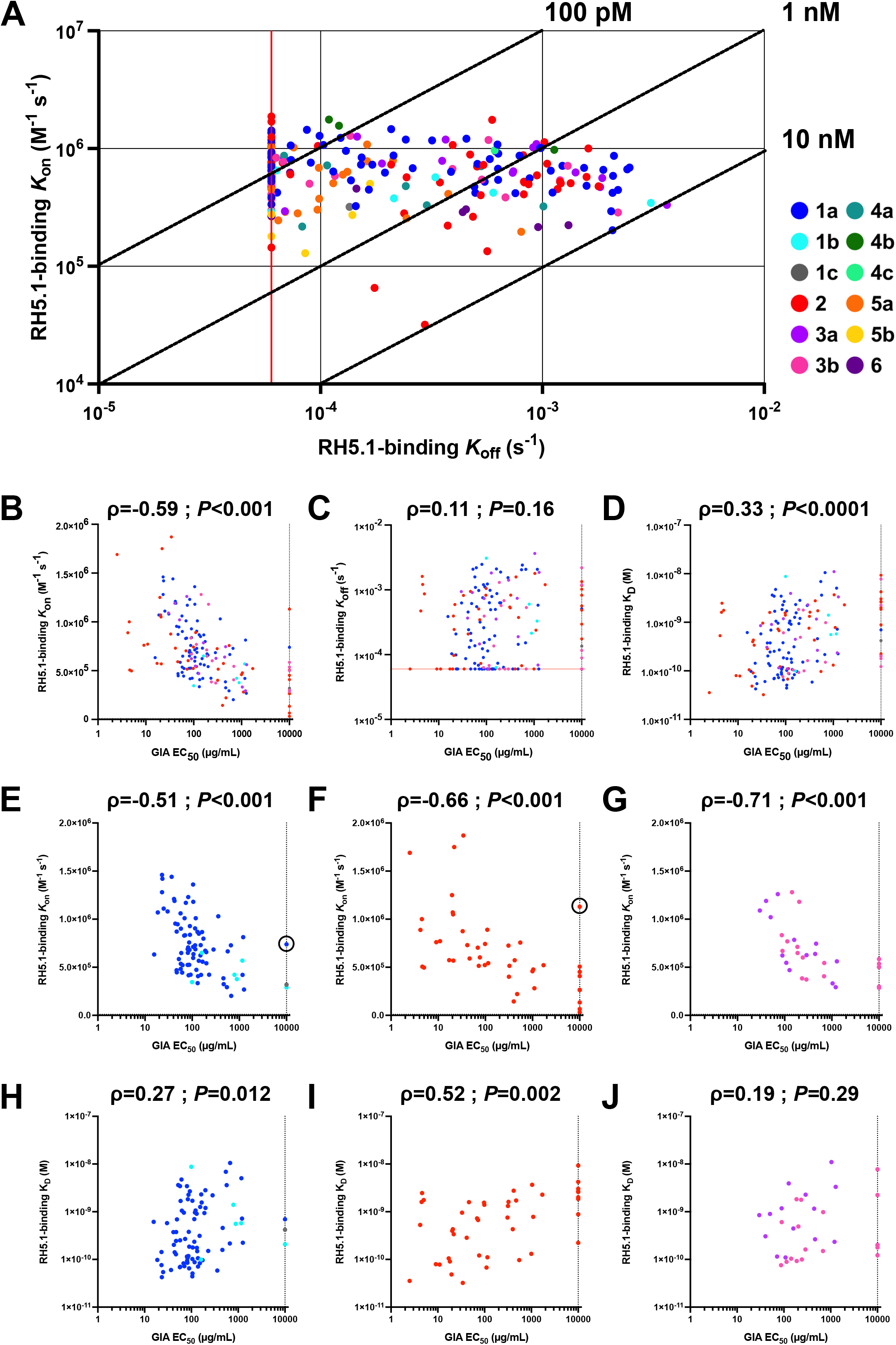
Binding kinetics of anti-PfRH5 mAbs. (**A**) Iso-affinity plot showing kinetic rate constants for binding of mAbs to RH5.1 (full-length PfRH5 protein) as determined by HT-SPR. Diagonal lines represent equal affinity (K_D_) = *K*_off_ / *K*_on_. Red vertical line indicates lowest limit of *K*_off_ measurement (6 x 10^-5^ s^-1^). mAbs colored by epitope community (N=213 in total). (**B**) The RH5.1-binding parameters of *K*_on_, (**C**) *K*_off_ and (**D**) K_D_ were correlated with GIA EC_50_ for all antibodies in the growth inhibitory antibody epitope (super)communities 1, 2 and 3 (N=159). (**E**) The RH5.1-binding parameter of *K*_on_ or (**H**) K_D_ was correlated with GIA EC_50_ for all antibodies in the growth inhibitory antibody epitope supercommunity 1 (N=83), (**F,I**) community 2 (N=44) and (**G,J**) supercommunity 3 (N=32). Anti-PfRH5 clones R5.129 and R5.036 are circled in panels **E** and **F**, respectively. Spearman’s rank correlation coefficient (ρ) and two-tailed *P* value are shown.

We next focused exclusively on community 2 and supercommunities 1 and 3 (together these contained nearly every growth inhibitory mAb in the entire panel) and assessed for correlation between the kinetic parameters and GIA EC_50_ as a measure of functional potency. The *K*_on_ and K_D_ parameters were highly correlated with antibody potency, whereas no correlation was observed with *K*_off_ (**Figure 2B-D**). The correlation between *K*_on_ and mAb GIA EC_50_ was also highly significant and comparable across all three (super)communities (**Figure 2E-G**). In contrast, the correlation between *K*_D_ and mAb GIA EC_50_ was weaker and only significant for (super)communities 1 and 2 (**Figure 2H-J**), and likely driven by the underlying correlation with *K*_on_. Two obvious exceptions to this trend were observed – R5.129 in community 1a and R5.036 in community 2 (circled, **Figure 2E-F**). These mAbs may bind to distal regions of their community’s epitope footprint on PfRH5; indeed, neither of these block basigin (**Figure 1B**), and both display competition interactions outside of their (super)community (**Figure 1A**). Overall, these data suggest speed of PfRH5 binding is a major determinant of growth inhibitory mAb potency, regardless of epitope binding site (within or close to the basigin binding site on PfRH5), and that a slow rate of binding is likely sufficient to render an antibody ineffective.

### Sequence analysis of anti-PfRH5 mAbs reveals a potent public clonotype

To complement the high-resolution epitope and functionality mapping of anti-PfRH5 antibodies, we next conducted a sequence analysis of all 206 mAbs. The variable heavy and light chain gene segment usage of each mAb was annotated using IMGT V-quest (**Data S3A**). The range of somatic hypermutation (SHM) in the variable heavy chain was comparable across the epitope communities, with a trend towards greater SHM in communities 3a, 4c, 5b and 6, and the converse in communities 1b, 4a and 4b (**Figure S3A**). Levels of SHM also differed by dosing regimen in the RH5.1/AS01_B_ clinical trial. Antibodies isolated from the delayed boosting groups showing significantly higher levels of SHM in their heavy and light chain gene segments as compared to the monthly boost group (**Figure S3B-C**); in line with these groups showing slower dissociation rates and improved affinity constants (**Figure S2C-D**). The median CDRH3 length of the N=206 mAbs was 14 amino acids, similar to the average length reported for the human IgG repertoire ^27^. However, individual epitope communities diverged from this median, with supercommunity 1 and community 6 using marginally shorter median CDRH3 lengths (**Figure S3D**). Antibodies with exceptionally long CDRH3 sequences, >20 amino acids, occurred in 8/12 communities. Communities 3a and 4b were noted for their bias towards these longer CDRH3 sequences, and had median lengths of 21.5 and 21 amino acids, respectively.

Analysis of heavy and light chain gene family usage across the whole anti-PfRH5 panel (**Figure 3A, Data S3B**) revealed a diverse repertoire of N=5 heavy chain and N=10 light chain gene families, although with a notable predominance of HV3 (N=85), HV4 (N=86), KV1 (N=88), KV3 (N=47) and LV3 (N=42) gene family usage. The HV4 gene family was used by most antibodies in community 1a (61/75, 81%), whereas the HV3 gene family was frequently used by community 2 (32/43, 74%), 3b (13/17, 76%) and 4a (10/10, 100%) antibodies. No single light chain gene family was used by more than 50 % of mAbs within a community, with the exception of LV3 which was used by 9/10 and 3/3 community 4a and 4b mAbs, respectively. Across these gene families, N=27 possible combinations of pairings were observed, with the HV4/KV3 pairing most frequently identified (N=38/206). Notably, some pairings were commonly associated with specific epitope communities – HV4/KV3 and HV4/KV1 in community 1a and HV3/LV3 in community 4a (**Figure S3E**), otherwise other common gene family pairings were often present in antibodies from different epitope communities. Finally, we analysed the pairings of individual genes, which similarly revealed a diverse repertoire; here the highest frequency gene pairing included only 8 mAbs (HV4-39/KV3-11) which were all community 1a antibodies, whilst 97/206 mAbs used a unique gene pairing (**Figure 3B, Data S3B**).

**Figure 3:**
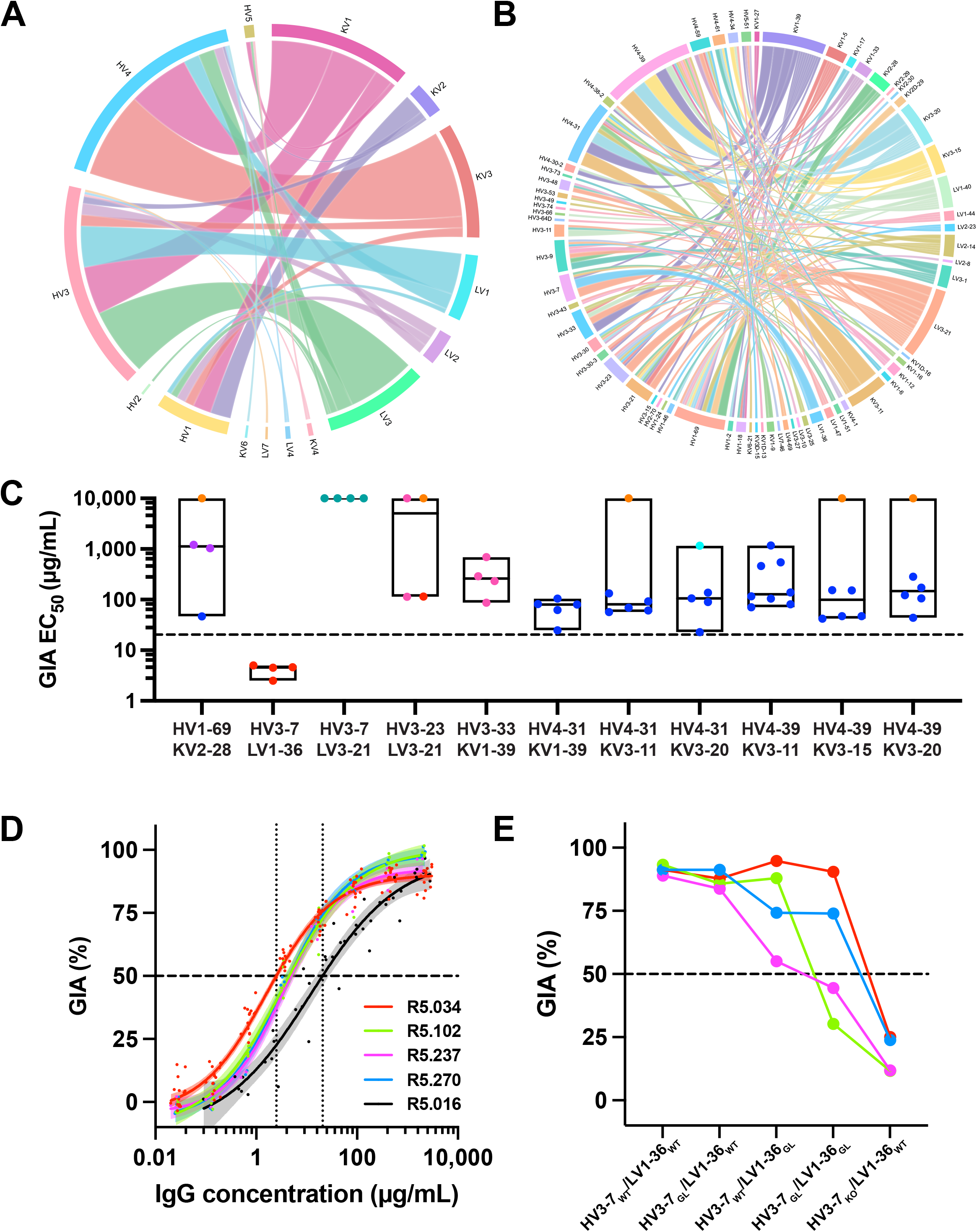
Sequence analysis of anti-PfRH5 mAbs. (**A**) Chord plot representing pairings of immunoglobulin HV and KV/LV gene families and (**B**) genes used by N=206 anti-PfRH5 human mAbs. The width of the cord is proportional to the number of mAbs which utilize that pairing. (**C**) Gene pairs in the anti-PfRH5 panel with N≥4 representative mAbs plotted in groups along with their GIA EC_50_ value. Each mAb is colored by its epitope community. Boxes show the mean with minimum to maximum. Dashed line shows the R5.016 bench mark EC_50_ of 20.7 µg/mL. (**D**) GIA assay titration curves of mAbs utilizing the HV3-7/LV1-36 gene combination and sentinel mAb R5.016 for comparison. Data were combined from repeat assays: N=14 for R5.034; N=4 for R5.237; N=3 for R5.237; N=3 for R5.270; and N=6 for R5.016. Data were log transformed and a four parameter nonlinear regression was fitted. Individual points are from the all replicate titrations. The shaded regions show the 95% confidence limits of the curves. Dashed line shown at 50 % GIA and dotted lines at EC_50_ readouts for the R5.034 and R5.016 curves. (**E**) For each mAb utilizing the HV3-7/LV1-36 public clonotype gene combination (color-coded as in **D**) a panel of four germline revertant antibodies was designed. Each mAb was tested at a concentration of 0.5 mg/mL in the GIA assay against 3D7 clone *P. falciparum* parasites. Points shown the mean of triplicate test wells and connecting lines are shown for clarity. WT = wild-type mAb sequence; HV3-7_GL_ has all mutations up to the beginning of the CDRH3 sequence mutated to germline combined with WT light chain; LV1-36_GL_ has all mutations reverted to germline, including the CDRL3 and J-region, combined with WT heavy chain; HV3-7_GL_/LV1-36_GL_ is mAb with both of these heavy and light chain sequences; HV3-7_KO_ is the HV3-7 WT sequence for each respective mAb but with the CDRH3 mutated to a random sequence of 13 amino acids, combined with WT light chain.

To assess for association of antibody gene pairing with mAb GIA EC_50_ potency, we initially analysed all gene pairs with N≥4 representative mAbs, which revealed a specific gene combination, HV3-7/LV1-36, of exceptional potency (**Figure 3C**). This same combination of heavy and light chain variable gene segment usage was also independently identified as predictive of high GIA by an unbiased computational modelling analysis of all available antibody gene sequence data across the mAb panel (**Figure S3F, Data S3C**). Notably all four of the HV3-7/LV1-36 mAbs were in community 2, were independently isolated from four separate vaccinees, had the same CDRH3 length (**Figure S3G**), and had GIA EC_50_s below 5 µg/mL up to 8-fold more potent than the previous best-in-class human mAb R5.016 ^24^ (**Figure 3D**). This grouping encompassed 4/5 of the highest potency mAbs identified across the entire panel, including the most potent clone R5.034 (**Figure 1D**) along with R5.102, R5.237 and R5.270; the single exception in this high potency cluster was mAb R5.268 which utilized HV3-48/LV3-21. We further explored the contribution of gene sequence to potency by producing a panel of germline revertants. Although SHM of the light chain gene contributed to GIA potency for some of the mAbs, all were highly dependent on their CDRH3s for mediating parasite growth inhibition (**Figure 3E**). The HV3-7/LV1-36 gene combination thus, in summary, defined a highly potent anti-PfRH5 public clonotype.

### Structural definition of the PfRH5 antigenic landscape

We next mapped the epitope (super)communities onto the structure of PfRH5. Structural information existed for sentinel mAbs across most super(communities), however, supercommunity 3 was undefined. To address this, we obtained a 3.3 Å structure of R5.251 (community 3a) in complex with RH5ΔNL (**Table S1**). This clone was the most potent identified in supercommunity 3, with a GIA EC_50_ of 29 µg/mL – similar to the previously reported best-in-class antibody R5.016 (community 2) and 3-fold more potent than the sentinel R5.008 (GIA EC_50_ 90 µg/mL, community 3b) ^24^. R5.251 bound around the tip of PfRH5 (**Figure 4A**). Analysis of the binding interface using PDBePISA predicted involvement of 18 residues on R5.251 (**Data S4**) primarily through the CDRH3, CDRL1 and CDRL3 loops, with no direct CDRH1 or CDRH2 involvement (**Figure 4B**). Docking of basigin to this structure showed steric clashes between basigin ^19^ and the light chain of R5.251 (**Figure 4C**), consistent with its ability to block basigin in the direct blocking assay (**Figure 1B**).

**Figure 4:**
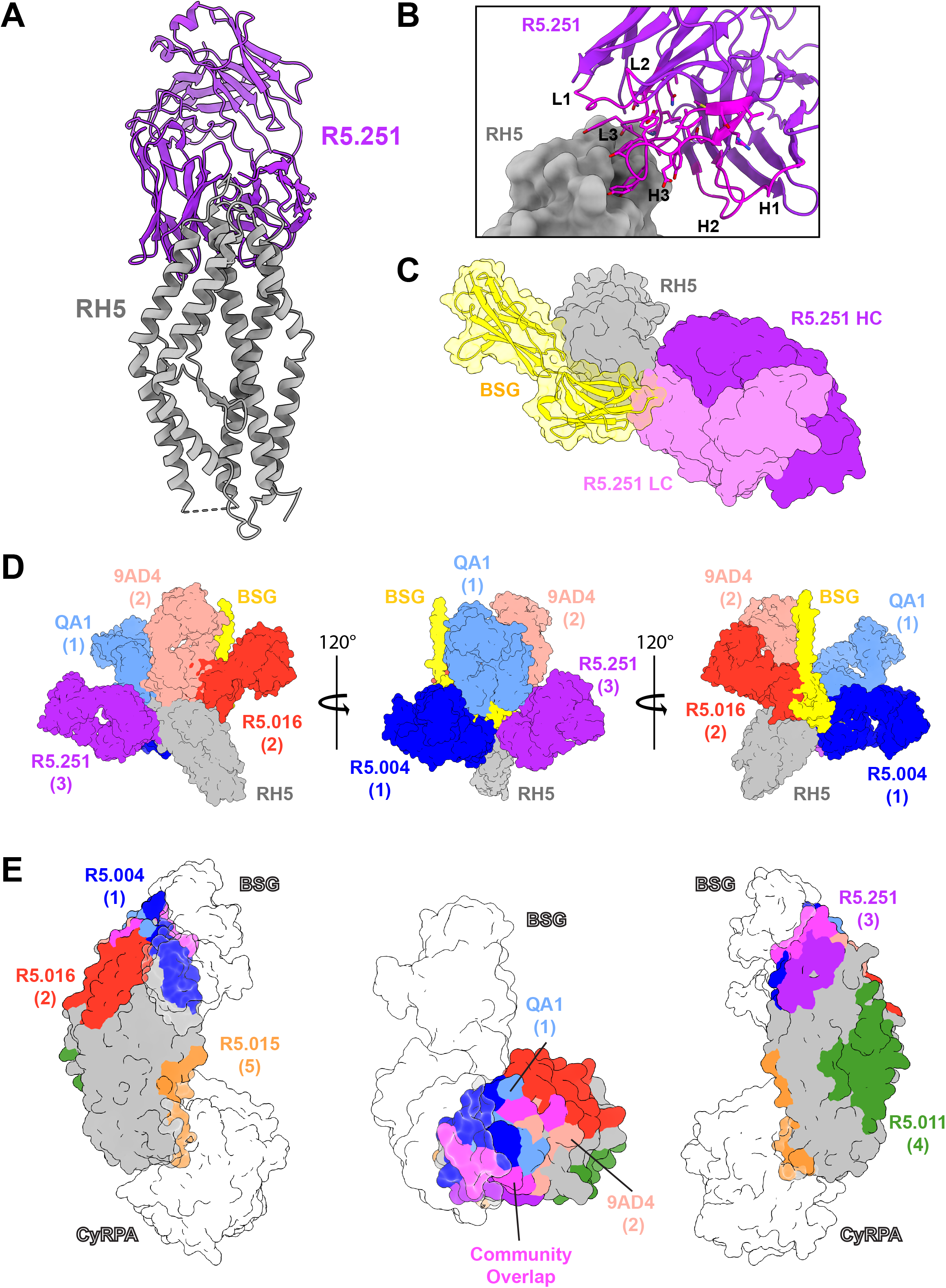
The antigenic landscape of PfRH5. (**A**) Crystal structure of PfRH5 using RH5ΔNL protein (grey) bound to R5.251 Fab fragment (violet). (**B**) Close-up view of the PfRH5 (grey) and R5.251 (violet) binding interface. Complementarity determining regions (CDRs) are highlighted in magenta, and labelled with their identifier, according to IMGT annotation. (**C**) Structure of human basigin (CD147, yellow) (PDB: 4UOQ) ^19^ aligned to the structure of PfRH5 (grey) in complex with the heavy (violet) and light (pink) chains of R5.251 Fab. (**D**) Structure of PfRH5 (grey) and R5.251 Fab (violet, community 3a) aligned to the structures of Fabs R5.004 (blue, community 1a, PDB: 6RCU); R5.016 (red, community 2, PDB: 6RCU) ^24^; QA1 (pale blue, community 1a, PDB: 4U1G) and 9AD4 (pale red, community 2, PDB: 4U0R) ^19^; and to basigin (yellow, PDB: 7CKR) ^28^. (**E**) Structure of PfRH5 (grey, PDB: 4WAT) ^29^ colored by the interface residues with Fabs R5.251 (violet, community 3a); R5.004 (blue, community 1a, PDB: 6RCU); R5.011 (green, community 4b, PDB: 6RCV) ^24^; R5.015 (orange, community 5b, PDB: 7PHU) ^25^; R5.016 (red, community 2, PDB: 6RCU); QA1 (pale blue, community 1a, PDB: 4U1G) and 9AD4 (pale red, community 2, PDB: 4U0R). Interfacing residues used by two or more different communities are highlighted in magenta. Basigin (PDB: 4U0R) ^19^ and CyRPA (PDB: 6MPV) ^20^ are shown as silhouettes. The leftmost and rightmost images are flipped 180° relative to one another. The centre images is a top-down view, centred on the apex of PfRH5; PfCyRPA has been omitted from this view.

We next docked into this model the basigin ectodomain and transmembrane helix ^28^ and the other available structures of growth inhibitory anti-PfRH5 human antibodies – sentinel mAbs R5.004 and R5.016 from (super)communities 1 and 2, respectively ^24^. We also supplemented these with the available structural data on two mouse mAb clones, QA1 and 9AD4 ^19^, which we placed into communities 1a and 2, respectively, using our new mAb panel (**Figure S4A**). These data revealed a “crown” of growth inhibitory mAbs, with varied footprints and binding angles, that decorate around the region of the basigin binding site on PfRH5 (**Figure 4D**). We next determined the interfacing residues of all anti-PfRH5 antibody clones on the surface of PfRH5 ^29^ further supplementing the above analysis with the available structural data on sentinel mAbs R5.011 and R5.015 from supercommunities 4 and 5, respectively ^24,25^ (**Figure S4B**). This provided a complete map of all five epitope (super)communities as recognized by these Fabs on the alpha-helical diamond core of PfRH5 (**Figure 4E**).

### Multiple intra-PfRH5 mAb interactions modulate parasite growth inhibition

Having characterized the functional, biophysical and structural properties of the mAb panel, we next sought to assess for functional interactions between the various epitope super(communities). We previously reported that a non-inhibitory antibody (R5.011, community 4b) was able to potentiate or synergize with the growth inhibitory sentinel antibodies R5.004, R5.016 and R5.008 (communities 1a, 2 and 3b, respectively) in the GIA assay ^24^. We thus systematically screened for this phenotype of functional intra-PfRH5 antibody interaction on a large scale and also sought to assess whether the high GIA EC_50_ potency of the HV3-7/LV1-36 public clonotype could be outperformed via a combination of anti-PfRH5 mAbs. Initially we devised a screening strategy to test growth inhibitory or “neutralizing” antibodies (nAbs) in combination with non-neutralizing antibodies (non-nAbs). Nine representative nAbs (of high potency wherever possible and spanning communities 1a, 1b, 2, 3a and 3b and the HV3-7/LV1-36 public clonotype) were selected; all blocked basigin in the BLI assay, with the exception of R5.034 and R5.102 in community 2. These were subsequently screened for GIA in pair-wise combinations with a further 23 non-nAbs that spanned all applicable epitope communities, i.e., those with at least one non-nAb (1a, 1c, 2, 3b, 5a, 5b and 6) and also all clones in supercommunity 4 (which includes R5.011). All nAbs were tested in the GIA assay at their EC_50_ concentration, with and without addition of each non-nAb held at 0.3 mg/mL (**Figure S5A**). The predicted Bliss additivity GIA value ^30,31^, calculated from the % GIA of the nAb and non-nAb tested alone, was subtracted from the % GIA of the test combination, with thresholds defined to categorize pairings as synergistic, additive, or antagonistic (**Figure 5A**).

**Figure 5:**
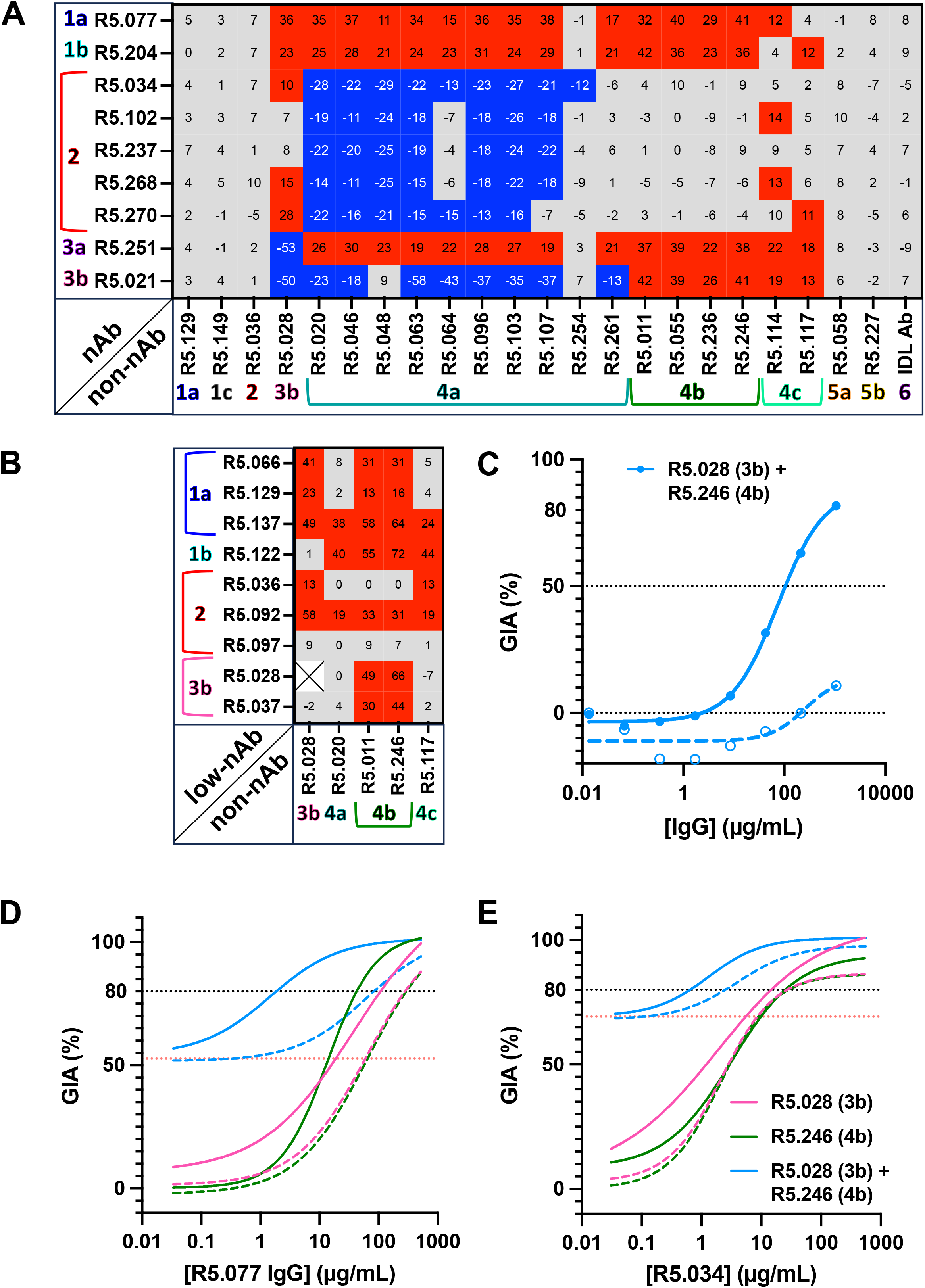
Assessment of intra-PfRH5 antibody synergy. (**A**) Growth-inhibitory/neutralizing antibodies (nAbs) were tested at a final concentration equivalent to their EC_50_ value and non-nAbs were tested at a final concentration of 0.3 mg/mL. The predicted Bliss additivity % GIA was subtracted from the measured % GIA of the test antibody combination, and the difference is plotted as a percentage in the heatmap. Thresholds were used to categorize combinations as synergistic (≥10%; red), additive (grey), or antagonistic (≥–10%; blue). Test mAbs are annotated with their epitope community assignment. (**B**) As for panel **A** except pairs of non-nAbs or mAbs with minimal GIA were combined. Antibodies were tested at a final concentration of 0.2 mg/mL each. (**C**) GIA assay dilution curve of R5.028 (community 3b) in combination with R5.246 (community 4b). Antibodies were combined in an equal ratio and run in a 5-fold dilution curve starting from ∼1 mg/mL. Data were log transformed and a four-parameter non-linear regression was plotted. The predicted Bliss additivity % GIA of the mixture is shown as a dashed curve. A dotted line is shown at 50 % GIA for reference. (**D**) GIA assay dilution curves of the community 1a nAb R5.077 run in a 5-fold dilution starting from ∼0.5 mg/mL under various test conditions. Data were log transformed and a four-parameter non-linear regression was plotted. For each curve, a non-nAb (R5.028 from community 3b or R5.246 from community 4b), or a combination of both non-nAbs was added at a fixed concentration of 0.2 mg/mL each. Predicted Bliss additivity GIA curves for each combination are shown as a dashed line. The black dotted line indicates 80 % GIA. The red dotted line indicates the level of GIA measured alone for the fixed concentration combination of the two non-nAbs (R5.028+R5.246). (**E**) Same assay set-up as in **D** except using R5.034 (community 2) in place of R5.077.

Non-nAbs from communities 1a, 1c, 2, 5a, 5b and 6 showed no obvious interactions with any nAb in the test panel, neither reducing nor potentiating the overall level of GIA. In contrast, the non-nAbs from community 4b (including R5.011) were consistently synergistic with mAbs from supercommunities 1 and 3, but did not affect the GIA of nAbs from community 2. Interestingly non-nAbs that comprise community 4a showed the same potentiating effect with nAbs from communities 1a, 1b and 3a. However, they were antagonistic with nAb clones from community 2 (despite no competitive binding for PfRH5) and from community 3b, although this was likely due to some competitive binding. Community 4c showed a less consistent profile, although overall these clones were able to synergize with some nAbs from each community, albeit weakly. Notably, the representative non-nAb from community 3b (R5.028) could synergize with nAbs from communities 1a, 1b and 2; whilst exhibiting antagonism with the two nAb clones from supercommunity 3 (R5.021 and R5.251) with which it competes for PfRH5 binding. Further testing using a range of concentrations of representative nAbs from communities 1a (R5.077), 2 (R5.034, R5.268) and 3a (R5.251) in combination with a fixed concentration of representative modulatory non-nAbs from communities 3b, 4a, 4b and 4c, confirmed these results (**Figure S5B-C**).

Having shown a range of non-nAb specificities could modulate the GIA of nAb clones, we also assessed whether the potentiating non-nAb clones from epitope communities 3b, 4a, 4b and 4c could synergize with antibody clones that exhibit very low or no GIA. Here we selected clones from communities 1a, 1b, 2 and 3b that originally showed poor GIA assay EC_50_ outcome (**Figure 1D**) in contrast to the majority of other nAbs in each of these communities. In this case, although a number of antibody combinations continued to show minimal or no GIA, over half of the pairs of clones tested now showed improved GIA and these spanned across the range of epitope communities (**Figure 5B**). Moreover, the non-nAbs from community 3b also showed synergy with the clones from community 4b, including the potentiating clone R5.028. Further analysis of the combination of R5.028 (community 3b) and R5.246 (community 4b) showed how these two GIA-negative and non-basigin blocking mAbs could combine to give high-level synergy resulting in growth inhibition of parasite invasion (EC_50_ 105 μg/mL) when tested in a 1:1 mixture (**Figure 5C**).

Having defined non-nAbs from two non-competing epitope communities (3b and 4b) that both potentiate nAbs and each other, we next tested them in triple combination with the best nAb from community 1a (R5.077) and the most potent representative of the public clonotype from community 2 (R5.034). In the case of R5.077, a 45-fold increase in EC_80_ potency was observed under the GIA assay test conditions when both R5.028 and R5.246 were added (1.9 μg/mL versus 83.4 μg/mL for predicted additivity). This was far greater than that observed when combining with R5.028 (2.6 fold-change) or R5.246 (6.9 fold-change) alone (**Figure 5D**), suggesting that the synergistic effect of either or both antibody clones is enhanced in the presence of the other. A similar observation was seen with R5.034, despite the fact that mAbs from community 2 do not synergize with R5.246 alone. Here, the combination of R5.028 and R5.246 with R5.034 produced a greater fold change in EC_80_ (3.9-fold) than the predicted additive effect of the triple combination under the GIA assay test conditions (**Figure 5E**). Finally, we also replicated this phenomenon with a pool of polyclonal anti-PfRH5 IgG from the RH5.1/AS01_B_ vaccine trial (**Figure S5D**), suggesting there remains substantial room for improvement in terms of the qualitative potency of the vaccine-induced antibody response. However, notably, most of these previous experiments involved testing a titrated concentration of one test antibody with another held at fixed concentration. We thus finally sought to identify the most potent anti-PfRH5 mAb or combination on a per µg basis for assessment as a novel blood-stage anti-malarial intervention. A series of single mAbs and combinations were down-selected based on the previous analysis of synergistic interactions. Each antibody mixture was compared head-to-head in a titration analysis in the assay of GIA against 3D7 clone *P. falciparum* parasites. Although some of these combinations could almost match R5.034 in potency, none could ultimately outperform the EC_50_ of the single most potent clone R5.034 from community 2 (**Figure S5E-G**).

### The public clonotype antibody R5.034 protects against *P. falciparum* sporozoite challenge

Given the R5.034 public clonotype mAb demonstrated the most potent GIA EC_50_ across all of our analyses, we investigated the merits of this clone as a prophylactic intervention against *P. falciparum*. To better understand the structural and binding characteristics of this candidate, we obtained a crystal structure of R5.034 (resolved to show its Fv region only) in complex with RH5ΔNL to 4 Å (**Figure 6A**, **Table S1**). Analysis of the binding interface using PDBePISA showed that R5.034 bound an upper facet, close to the tip of PfRH5. Within this interface, only 5/13 residues used by the heavy and light chains of R5.034 (excluding those in the CDRH3) were mutated from germline HV3-7/LV1-36 sequence (**Data S4**), in line with this mAb’s tolerance of germline reversion mutation outside the CDRH3 (**Figure 3E**). In the case of PfRH5, the binding interface was centred on a 3-helical bundle of the PfRH5-fold, with the CDRH3 loop projecting towards a cleft created by the outermost two α-helices. The interfacing area on PfRH5 spanned 19 amino acids (**Data S4**) and contained none of the few commonly observed polymorphisms ^32^, suggesting that the epitope of R5.034 is conserved. Comparison of R5.034 with the other structurally characterized mAbs from community 2 (R5.016 and 9AD4) demonstrated that R5.034 shared a similar overlapping epitope as predicted by the competition data. All three antibodies bound close to the basigin binding site, with their Fab constant regions (modelled for R5.034) projecting into the space predicted to be occupied by the erythrocyte membrane (**Figure 6B**).

**Figure 6:**
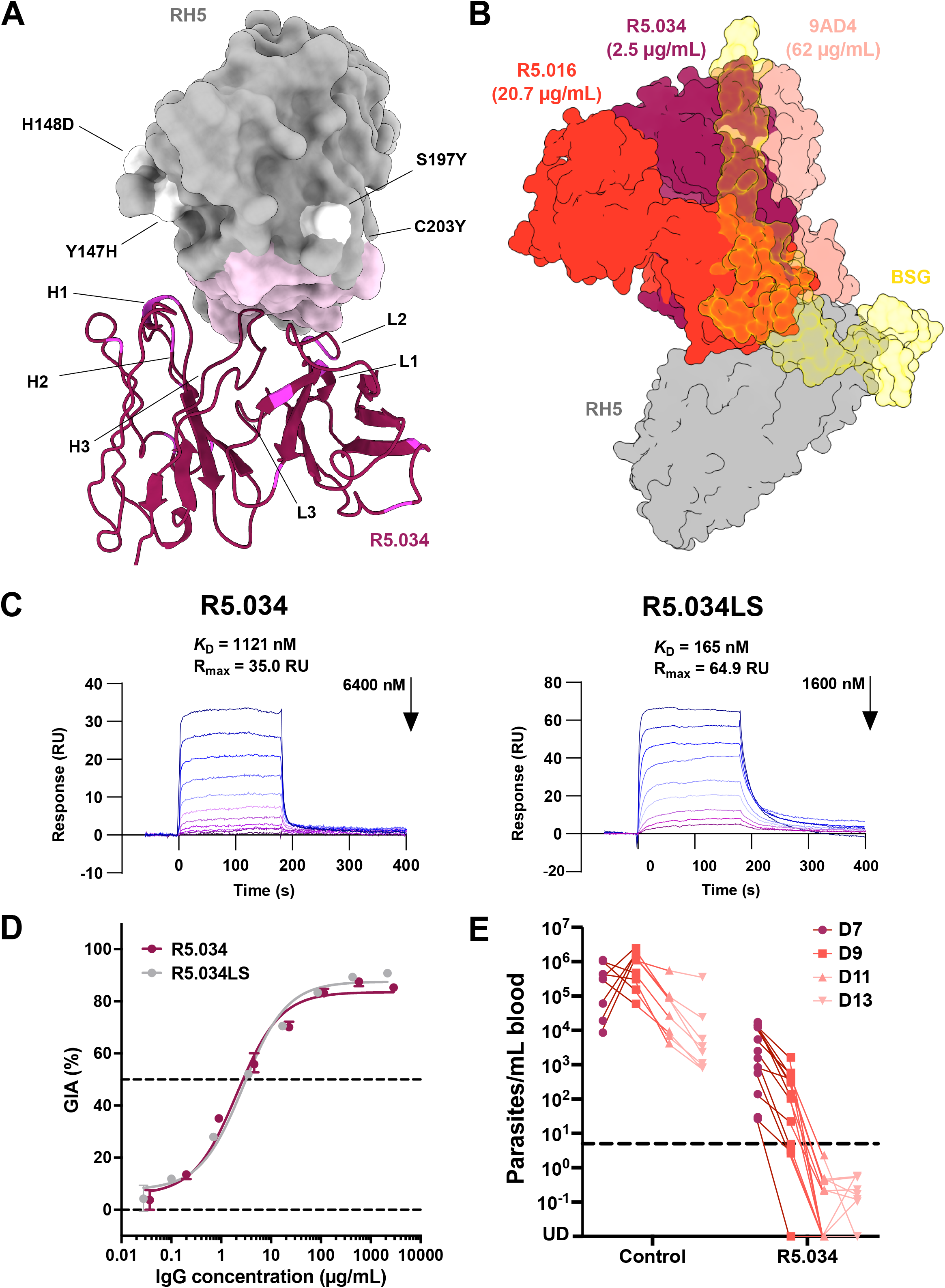
Structure of R5.034 and efficacy against *P. falciparum* challenge. (**A**) Crystal structure of PfRH5 using RH5ΔNL protein (grey) in complex with R5.034 Fv region (maroon). The image is of a top-down view, tilted 10° along the x-axis to view the binding interface on PfRH5 as predicted by PDBePISA, which is coloured in pink. Common PfRH5 polymorphisms are colored white and annotated. CDR loops in the R5.034 structure are labelled with their IMGT identifier. Residues in the HV and LV regions which are mutated from germline are highlighted in magenta. (**B**) Structure of PfRH5 (grey) and R5.034 modelled Fab (maroon) aligned with the community 2 Fabs of 9AD4 (pale red, PDB: 4U0R) ^19^ and R5.016 (red, PDB: 6RCU) ^24^ and the basigin ectodomain with transmembrane helix (yellow, PDB: 7CKR) ^28^. An AlphaFold predictive model ^40,41^ of R5.034 Fab was aligned with the experimentally observed R5.034 Fv structure to hypothesize the spatial arrangement of the three antibodies. (**C**) Steady-state affinity, as assessed using SPR, of the R5.034 and R5.034LS mAb binding to human FcRn at pH 6.0. Sensorgrams are shown of a 9-step dilution curve beginning at the indicated concentration of mAb. Calculation of steady-state affinity (K_D_) at pH 6.0 is shown in **Figure S6B**. (**D**) Titration of the R5.034 and R5.034LS mAbs in the assay of GIA against 3D7 clone *P. falciparum* parasites. Dots show the mean and error bars the range of N=3 triplicate test wells per test mAb concentration. Non-linear regression curve is shown. (**E**) FRG huHep mice were each exposed to the equivalent of five bites using mosquitos infected with N54 strain *P. falciparum*. On day 5 post-infection, mice were administered 675 µg control mAb or 625 µg R5.034, as well as human RBC, via the intravenous route. Administration of more human RBC was repeated on days 6, 9 and 11. Parasitemia in the blood was monitored on days 7, 9, 11 and 13 by quantitative RT-PCR. Data from individual mice are shown combined from two independent experiments (Control N=8 ; R5.034-treated N=13). Dashed line is the lower limit of detection of the qRT-PCR assay at 5 parasites per mL blood.

We subsequently generated a second version of this antibody with the “LS” mutation in the IgG1 Fc domain (R5.034LS) used in the clinical development of candidates mAbs to extend serum half-life via increased antibody recirculation ^33^. R5.034 and R5.034LS demonstrated comparable binding kinetic rate constants and high affinities for PfRH5 binding of 30-40 pM (**Figure S6A**). To further assess the LS half-life extension mutation, we determined binding to the human neonatal Fc receptor (FcRn). Here, as expected, neither R5.034 or R5.034LS bound to FcRn at pH7.4 (**Figure S6B**), but at lower pH6.0 (to mirror endosomal recycling of plasma IgG), R5.034LS showed an ∼7-fold higher affinity for FcRn as compared to the wild-type R5.034 IgG1 molecule (**Figure 6C, S6B**). Further screening in the GIA assay confirmed both R5.034 and R5.034LS exhibited identical functional potencies against *P. falciparum in vitro* (**Figure 6D**). Finally, we assessed protective efficacy of R5.034 passive transfer prior to *P. falciparum* mosquito-bite challenge in a humanized mouse model (**Figure 6E**). Control animals developed very high-level blood-stage parasitemia following parasite emergence from the liver, starting from a median level of 3.2×10^5^ parasites per mL of blood (p/mL) on day 7 and peaking on day 9 at 7.9×10^5^ p/mL. This declined over time, but all animals remained parasitemic on day 13 when the experiment was ended. In contrast, animals receiving R5.034 peaked on day 7 at a median level of 2.5×10^3^ p/mL (100-fold lower than controls) with this difference widening by day 9, whereby all animals showed decreased parasite burden to a median level of 1.4×10^2^ p/mL (5600-fold lower than controls). All animals receiving R5.034 were subsequently parasite negative, as measured by qRT-PCR, on day 11. Serum antibody levels in the R5.034-treated animals reached a median of 93 µg/mL (range: 48-98 µg/mL) on day 6 and were maintained at 83 µg/mL (range: 30-122 µg/mL) on day 13 (**Figure S6C**). At these antibody levels, R5.034 thus showed high-level efficacy against blood-stage *P. falciparum*.

## DISCUSSION

Here we provide the first high-resolution analysis of the antigenic landscape of the most advanced blood-stage malaria vaccine candidate antigen, PfRH5, and characterize in detail the biophysical, structural and combinatorial features of the human IgG antibody response that associate with functional blood-stage growth inhibition of *P. falciparum*. In doing so, we also identified an anti-PfRH5 public clonotype that shows the highest levels of *in vitro* growth inhibition reported to-date and confirmed its prophylactic ability to control and clear high-level blood-stage *P. falciparum* parasitemia in humanized mice. To perform our analyses, we isolated and characterized over 200 anti-PfRH5 mAbs from healthy UK adults immunized with the full-length protein-in-adjuvant vaccine RH5.1/AS01_B_ ^13,16^; the RH5.1 vaccine has since entered Phase 2b field efficacy testing in infants in Burkina Faso reformulated in Matrix-M™ adjuvant (ClinicalTrials.gov NCT05790889). Previous analyses of RH5.1/AS01_B_ vaccinee sera have shown the disordered N- and C-termini of the PfRH5 molecule do not contribute growth-inhibitory antibodies, in line with recent data showing that N-terminal cleavage of PfRH5 occurs in the micronemes prior to movement of PfRH5 to the merozoite’s surface ^34^. We therefore focussed our analysis on the structured core of the PfRH5 molecule (“RH5ΔNC”; amino acids K140-N506) ^19^, where we determined antigenic sites in line with the network of competitive binding interactions across the mAb panel. We thus defined 12 communities of human antibodies that bind PfRH5, and 10 of these clustered into four supercommunities.

Over 95 % of the mAb panel recognized conformational epitopes; of the few that could bind denatured RH5.1 protein, most bound to linear peptide epitopes within the IDL region (amino acids N248-M296) and we grouped these mAbs into community 6. We could not identify any functional outcomes for these antibodies; they neither mediated parasite growth inhibition nor showed any detectable interaction with PfRH5 binding partners or other anti-PfRH5 mAbs. Serological analyses of the anti-PfRH5 polyclonal IgG response in human vaccinees suggest responses against the IDL are common ^17^; these data would suggest this antigenic region should be removed in a next-generation PfRH5 vaccine immunogen aiming to improve the qualitative potency of vaccine-induced antibody responses.

PfRH5 is delivered to the merozoite surface at the end of an elongated hetero-pentameric invasion complex, within which it forms an interaction with PfCyRPA ^9,20^. Antibodies capable of blocking the PfRH5-PfCyRPA interaction were exclusively contained within supercommunity 5. Conflicting reports in the literature, using individual or very small panels of antibodies ^23–25,35^, have debated whether blockade of the PfRH5-PfCyRPA interaction has functional consequence for inhibition of parasite invasion. Here, like for community 6 antibodies, we could ascribe no other functional outcomes to these clones, neither with regard to *in vitro* parasite growth inhibition nor modulatory interactions with anti-PfRH5 mAbs from other communities. Our analysis of ∼30 mAbs thus strongly suggests that blockade of this interaction cannot block parasite invasion of the RBC, and that the antigenic surface of PfRH5 ascribed to supercommunity 5 is covered by PfCyRPA and therefore buried within the invasion complex at the point of exposure to the host immune system.

Notably, our data show growth inhibitory anti-PfRH5 antibodies span five epitope communities: 1a, 1b, 2, 3a and 3b. Although structural data existed for exemplar antibodies spanning communities 1a, 1b and 2 ^19,24^, there were none from supercommunity 3. We therefore resolved the structure of mAb R5.251, the best-in-class from this supercommunity. Mapping of these structural data onto the α-helical core of PfRH5 shows the footprints of these epitope communities form a “crown” on the top half of the diamond-like architecture, overlapping with or adjacent to the basigin-bind site. Communities 1a, 1b and 3a were composed of mAbs that i) blocked basigin-binding as assessed by BLI, and ii) with only one exception showed functional GIA. Combined these accounted for almost half of the antibodies analysed in this study. In contrast, community 2 antibodies (the second largest cluster in the panel after community 1a) bind close or adjacent to the PfRH5-basigin interaction site, explaining why ∼70% of these mAbs failed to block basigin binding as measured here by BLI. Notably, the range of GIA potency in this community was the largest (ranging from GIA negative to the most potent clones identified) and GIA positivity did not associate with measurement of direct basigin-binding blockade. Other data now suggest basigin is present within macromolecular complexes in the RBC membrane, and that community 2 antibodies may in fact inhibit growth by sterically blocking the interaction of PfRH5 with basigin in its physiological context ^26^. Whether this kind of blocking assay could better predict GIA positivity across a large panel of community 2 mAb clones remains to be determined. In contrast, a key finding from our analysis was the highly significant correlation between mAb potency in the GIA assay and the association rate (*K*_on_) of PfRH5 binding but not the dissociation rate (*K*_off_), in line with the theorized kinetic constraints of an anti-merozoite vaccine ^36^. Notably, this relationship transcended (super)communities 1, 2 and 3, suggesting the speed with which an antibody engages PfRH5 at any of these three major antigenic sites is a central driver of its growth inhibitory potency against the rapidly invading *P. falciparum* merozoite. Indeed, this analysis builds on previous data reported for small panels of mAbs targeting both *P. falciparum* and *P. vivax* merozoites ^21,24,37^, and now strongly suggests that combining these antibodies with other anti-malarial antibodies or drugs that slow merozoite invasion could offer novel strategies to improve functional potency of anti-PfRH5 IgG.

The community 3b epitope region is next to the site of supercommunity 4, which itself descends down one side of the PfRH5 diamond-like structure, on the opposite side to the PfCyRPA-binding site recognized by mAbs in supercommunity 5. Over half of the community 3b mAbs blocked basigin-binding as assessed by BLI and showed a similar range of GIA potencies to community 1b, whilst the remaining clones in community 3b and all of those in supercommunity 4 were otherwise GIA negative. However, we previously reported mAb R5.011 (the sentinel clone for community 4b) was able to potentiate or synergize with growth inhibitory antibodies in the GIA assay, via a 3-fold lengthening of *P. falciparum* merozoite invasion time ^24^. Our analysis with this much larger panel of mAbs now defines this phenomenon in far greater resolution, indicating that multiple intra-PfRH5 mAb interactions can modulate anti-parasitic growth inhibition. We found one main example of consistent antagonism in the GIA assay between antibodies from community 4a and those from community 2; this warrants further investigation given the apparent physical separation of their binding sites on PfRH5 and lack of competitive binding. Notably, otherwise, our data show the antigenic sites spanning community 3b and all of supercommunity 4 can reproducibly elicit human antibody clones with the potentiating phenotype. Here, when combined, these GIA-negative clones synergize with or potentiate the basigin-blocking and growth inhibitory antibodies from communities 1a, 1b, 3a and 3b. Moreover, we provide the first evidence that two non-neutralizing and non-basigin-blocking anti-PfRH5 mAbs can combine to give high-level synergy resulting in GIA, as exemplified with the combination of clones from communities 3b and 4b. These data have important implications for how intra-PfRH5 antibody interactions are likely occurring within polyclonal anti-PfRH5 IgG to inhibit growth of *P. falciparum* parasites, and indeed offer a possible explanation for the impressive qualitative potency (i.e., low GIA EC_50_) of vaccine-induced PfRH5-specific antibody reported across the Phase 1 vaccine trials ^18^. Further studies are now underway to investigate the composition of the plasma anti-PfRH5 polyclonal IgG response in human vaccinees with respect to these phenomena.

These anti-PfRH5 antibody data also provide the high-resolution insight required to guide the rational design of next-generation PfRH5 vaccine immunogens that seek to induce responses that are qualitatively superior to the current clinical vaccine RH5.1. Indeed these results suggest focusing on immunogen designs that incorporate (super)communities 1-3, 4b and 4c with removal or masking of antagonizing community 4a and non-functional (super)communities 5-6. Importantly, antibodies recognizing these functional epitope sites can arise from a diverse range of human antibody gene usage, however, human vaccine delivery technologies, adjuvants and/or regimens that can also drive improved *K*_on_ rates need to be investigated. Previous serological data from the RH5.1/AS01_B_ vaccine trial suggested that delayed, as opposed to monthly, vaccine boosting in humans could improve memory B cell responses as well as serum antibody response longevity and avidity; however, the purified total IgG with more avid anti-PfRH5 IgG failed to show improved GIA potency ^13,38^. Our data now explain this result, because although the antibody clones isolated from the delayed booster vaccinees showed more SHM and higher RH5.1 binding affinity (K_D_), the underlying driver for this was significantly slower dissociation rates, as opposed to faster association rates, which we would not predict to improve GIA.

Finally, our data identified that an exceptionally potent class of anti-PfRH5 antibody can derive from the antibody gene combination HV3-7/LV1-36. This public clonotype identified a small subset of antibodies within community 2, all with GIA EC_50_s below 5 µg/mL against 3D7 clone *P. falciparum*. Structural and mutational analyses of the most potent clone, R5.034, showed PfRH5 binding at this conserved epitope is driven by its CDRH3. Moreover, passive transfer of R5.034 into humanized mice challenged with *P. falciparum* sporozoites showed reductions in blood-stage parasitemia, as compared to controls, in the range of 2-3 orders of magnitude following parasite emergence from the liver, and then absence of detectable parasites by qRT-PCR within 4 days whilst all controls remained parasitemic. Given the recent exciting clinical advances with the L9LS and TB31F candidate mAbs against the sporozoite- and transmission-stages of *P. falciparum*, respectively ^6,39^, the data here identify a new blood-stage mAb candidate, R5.034LS, that could be considered for clinical development as part of a multi-stage multi-mAb approach to achieve high-level single-shot prophylaxis against *P. falciparum* malaria.

## ACKNOWLEDGEMENTS

The authors are grateful for the assistance of David Pulido, Daniel Alanine, Robert Ragotte, Geneviève Labbé, Julie Furze, Fay Nugent, Wendy Crocker, Charlotte Hague, Jenny Bryant, Lana Strmecki, Andrew Worth (University of Oxford); Sally Pelling-Deeves for arranging contracts (University of Oxford); Robert Hedley and Vasiliki Tsioligka for assistance with flow cytometry (Sir William Dunn School of Pathology, University of Oxford); Angela Lee and Hubert Slawinski for support with 10X BCR sequencing (Oxford Genomics Centre); Diamond Light Source for beamtime (proposal mx28172) and the staff of beamlines I03, I04 and I24 for assistance with crystal testing and data collection; Ken Tucker and Timothy Phares (Leidos); Robin Miller (USAID); and all the VAC063 trial participants.

This work, as well as the VAC063 clinical trial, was made possible in part through support provided by the United States Agency for International Development (USAID) Malaria Vaccine Development Program (MVDP) (AID-OAA-C-15-00071 and 7200AA20C00017). The findings and conclusions are those of the authors and do not necessarily represent the official position of USAID. This work was also supported in part by the European Union’s Horizon 2020 research and innovation programme under a grant agreement for OptiMalVax (733273); the National Institute for Health Research (NIHR) Oxford Biomedical Research Centre (BRC) and NHS Blood & Transplant (NHSBT; who provided material), the views expressed are those of the authors and not necessarily those of the NIHR or the Department of Health and Social Care or NHSBT; the Bill and Melinda Gates Foundation (INV-005170 and INV-027520); and the University of Oxford Human Immune Discovery Initiative (HIDI) Internal Fund (0010370). BGW held a UK Medical Research Council (MRC) PhD Studentship [MR/N013468/1]; AJRC, LTW, KMi, CAL and JT are supported by the Division of Intramural Research of the National Institute of Allergy and Infectious Diseases (NIAID), National Institutes of Health (NIH); CMN held a Sir Henry Wellcome Postdoctoral Fellowship [209200/Z/17/Z], and SJD is a Jenner Investigator and held a Wellcome Trust Senior Fellowship [106917/Z/15/Z].

## AUTHOR CONTRIBUTIONS

Conceived and performed experiments and/or analysed the data: JRB, DP, NDW, AJRC, GG, DQ, AML, HD, CR, MA, BGW, WJB, NGP, TM, PK, LP, CDW, FRD, LDWK, LTW, JFP, SES, JdRS, BKW, LK, JT, CMN, KMc, SJD. Performed project management: KS, VK, ARN, RSM, CRK, AJB, LAS. Contributed reagents, materials, and analysis tools: AJRC, AMM, DAL, KMi, CAL, JT. Wrote the paper: JRB, KMc, SJD.

## INCLUSION AND DIVERSITY

We support inclusive, diverse, and equitable conduct of research.

## METHODS

### CONTACT FOR REAGENT AND RESOURCE SHARING

Further information and requests for resources should be directed to and will be fulfilled by the Lead Contact, Simon J. Draper.

#### Data and Code Availability

- Requests for monoclonal antibodies (mAbs) generated in the study should be directed to the Lead Contact, Simon J. Draper.
- Crystal structures of RH5ΔNL bound to R5.034 and R5.251 are deposited into the Protein Data Bank (PDB) under ID codes: 8QKS and 8QKR, respectively.
- The R markdown file for the prediction of GIA from germline gene usage (**Figure S3F**) has been included as a supplemental file (see **Data S3C**).

### EXPERIMENTAL MODEL AND SUBJECT DETAILS

#### Human Blood Sample Collection

All mAbs were obtained from samples of adult volunteers immunized with the RH5.1/AS01_B_ vaccine as part of the VAC063 clinical trial ^13^. VAC063 was an open-label, multi-center, dose-finding Phase I/IIa trial, including a controlled human malaria infection (CHMI) component, to assess the safety, immunogenicity and efficacy of the candidate *P. falciparum* blood-stage malaria vaccine RH5.1/AS01_B_. Volunteers were healthy, malaria-naïve UK adults ranging from 18-45 years of age. The study was conducted in the UK at the Centre for Clinical Vaccinology and Tropical Medicine, University of Oxford, Oxford, Guys and St Thomas’ NIHR CRF, London and the NIHR Wellcome Trust Clinical Research Facility in Southampton. The study received ethical approval from the UK NHS Research Ethics Service (Oxfordshire Research Ethics Committee A, Ref 16/SC/0345), and was approved by the UK Medicines and Healthcare products Regulatory Agency (Ref 21584/0362/001-0011). Volunteers signed written consent forms and consent was verified before each vaccination. The trial was registered on ClinicalTrials.gov (NCT02927145) and was conducted according to the principles of the current revision of the Declaration of Helsinki 2008 and in full conformity with the ICH guidelines for Good Clinical Practice (GCP).

Donations of human RBC from healthy adult volunteers for use in assays received ethical approval from the UK NHS Research Ethics Service (London – City & East Research Ethics Committee, Ref 18/LO/0415).

The RH5.1 vaccine was based on the full-length PfRH5 antigen (amino acids E26 - Q526) with 3D7 clone *P. falciparum* sequence, as reported in detail elsewhere ^16^. In brief, the vaccine was manufactured as a secreted soluble product from a stable *Drosophila melanogaster* Schneider 2 (S2) cell line and affinity purified via a C-terminal four amino acid (E-P-E-A) “C-tag” ^42^. All four putative N-linked glycosylation sequons (N-X-S/T) were mutated Thr to Ala. Volunteer samples from VAC063 Groups 1, 2, 3, 5 and 7 were used in this study ^13^. Participants in Groups 1, 2 and 5 received three “monthly” vaccinations of RH5.1/AS01_B_ at days 0, 28 and 56, with a dose escalation of RH5.1 from 2 μg (Group 1), to 10 μg (Groups 2 and 5). Volunteers in Group 3 received two 50 μg doses of RH5.1 followed by a final dose of 10 μg RH5.1 given at day 182 (a “delayed fractional dosing” (DFx) regimen). All vaccines were formulated in in 0.5 mL AS01_B_ adjuvant (GSK) regardless of RH5.1 protein dose. Group 5 underwent blood-stage CHMI with 3D7 clone *P. falciparum* 14 days after their third vaccination. Group 7 was composed of a subset of Group 5 vaccinees who went on to receive a final, delayed and fourth booster vaccination (D4thB) with 10 µg RH5.1/AS01_B_ approximately four months after their third vaccination followed by a second round of CHMI. PBMC samples were isolated and cryopreserved at approximately 2-4 weeks post-final vaccination for each group. Human blood samples were collected into lithium heparin-treated vacutainer blood collection systems (Becton Dickinson). PBMC were isolated and used within 6 h in fresh assays, otherwise excess cells were frozen in fetal calf serum (FCS) containing 10 % dimethyl sulfoxide and stored in liquid nitrogen. Plasma samples were stored at −80 °C. For serum preparation, untreated blood samples were stored at room temperature (RT) and then the clotted blood was centrifuged for 5 min at 1800 rpm. Serum was stored at −80°C.

#### Experimental Animal Models

The study using liver-humanized mice was carried out at the Oregon Health and Sciences University (OHSU) which is accredited by the Association for Assessment and Accreditation of Laboratory Animal Care International (AAALACi) and is a Category I facility with an approved Assurance (#A3304-01) on file with the Office for Laboratory Animal Welfare (OLAW), NIH, USA. The protocol was approved by the OHSU Institutional Animal Care and Use Committee (IACUC) under protocol IP00002077. FRG huHep mice on the NOD background were purchased from Yecuris, Inc. (Beaverton, OR, USA).

#### Cell Lines

Expi293F HEK cells were cultured in suspension in Expi293 expression medium (Thermo Fisher Scientific) at 37°C, 8 % CO_2_, on an orbital shaker set at 125 RPM. *Drosophila* S2 cells were cultured in suspension in EX-CELL 420 medium (Sigma-Aldrich) supplemented with 100 U/mL penicillin, 0.1 mg/mL streptomycin and 10 % foetal bovine serum (FBS) at 25 °C. Stable S2 cell lines expressing PfRH5 proteins were generated using the ExpreS^2^ platform (ExpreS^2^ion Biotechnologies) as previously described ^16,43^.

### METHOD DETAILS

#### Isolation of PfRH5-specific B cells

PfRH5-specific B cells were single cell sorted from the majority of cryopreserved PBMC samples as previously described ^38^ but with minor modification as follows. In brief, samples were thawed into R10 media (RPMI [R0883, MilliporeSigma] supplemented with 10 % heat-inactivated FCS [60923, Biosera], 100 U/mL penicillin / 0.1 mg/mL streptomycin [P0781, MilliporeSigma], 2 mM L-glutamine [G7513, Millipore Sigma]) and were then washed and rested in R10 for 1 h. B cells were enriched (Human Pan-B cell Enrichment Kit [19554, StemCell Technologies]) and then stained with viability dye FVS780 (565388, BD Biosciences). Next, B cells were stained with anti-human CD19-BV786 (Clone: SJ25C1, 563325, BD Biosciences) and anti-human IgG-BB515 (Clone: G18-145, 564581, BD Biosciences), as well as two fluorophore-conjugated PfRH5 probes. To prepare the probes, monobiotinylated PfRH5 was produced by transient co-transfection of HEK293F cells with a plasmid encoding BirA biotin ligase and a plasmid encoding a modified PfRH5. The PfRH5 plasmid was based on ‘RH5-bio’ (a gift from Gavin Wright; University of York, York, United Kingdom; Addgene plasmid #47780) ^44^. RH5-bio was modified prior to transfection to incorporate a C-terminal C-tag for subsequent protein purification, as well as a 15 amino acid deletion to remove the disordered C-terminus of PfRH5 and a 115 amino acid deletion from the linear N-terminus to produce a protein known as “RH5ΔNC” ^38,45^. Probes were freshly prepared for each experiment by incubation of mono-biotinylated RH5ΔNC with streptavidin-PE (S866, Invitrogen) or streptavidin-APC (405207, eBioscience) at an approximately 4:1 molar ratio to facilitate tetramer generation and subsequent centrifugation to remove aggregates (13,000–14,000 rpm [max microcentrifuge speed] at RT for 10 min). Following surface staining, cells were washed and kept on ice until acquisition on the MoFlo (Dako cytomation). RH5ΔNC-specific B cells were identified as live CD19+ IgG+ RH5ΔNC-APC+ RH5ΔNC-PE+ cells and single cell sorted into 96-well plates containing 10µL/well lysis buffer (10mM Tris [T3038, Merck], 1 unit/mL RNasin Ribonuclease Inhibitor [N2515, Promega]) and frozen at -80 °C.

A subset of eight mAbs (R5.242, R5.243, R5.244, R5.246, R5.247, R5.248, R5.249 and R5.250) were derived from a cell sort that included an additional cell hashing step (TotalSeq-C0251/C0252/C0253/C0254/C0255 anti-human Hashtag 1/2/3/4/5 antibodies, cat # 394661/394663/394665/394667/394669, Biolegend) prior to B cell enrichment and staining. These hashed samples were then either single-cell sorted into 96-well plates (as above), or pooled for B cell receptor (BCR) sequencing analysis using the 10X chromium platform; performed at the Oxford Genomics Center, University of Oxford, UK.

#### Monoclonal Antibody Cloning

Reverse transcription and nested PCR of antibody heavy (VH) and light (VL) chains was carried out as previously described and using previously reported primers ^24,46^. PCR products were purified using a PCR purification kit (Qiagen, 28006) and then cloned into the AbVec expression plasmids to produce recombinant human IgG1 mAbs (a gift from Patrick C. Wilson, University of Chicago, USA). In brief, plasmids and PCR products were 5’ digested using BshTI and 3’digested using SalI (AbVec-HCIgG1), XhoI (AbVec-LLC) and Pfl23II (AbVec-KLC) before ligation using QuickLigase (NEB). Ligation products were then used to transform MultiShot^TM^ StripWell TOP10 chemically competent *Escherichia coli* (Life Technologies) following the manufacturer’s instructions. Transformed cells were plated on LB agar 8-well Petri plates (Teknova) containing 100 µg/mL carbenicillin and grown at 37 °C overnight in a static incubator. Colonies were picked and sent to Source BioScience for sequencing, and those with productive antibody VH and VL sequences (analysed with Geneious® software) were inoculated and plasmids extracted using a QIAgen MINIprep kit. Sequence analysis was carried out using Geneious Prime®. Germline identity and gene usage parameters were determined using IMGT/V-QUEST. Heavy chain CDR3 (CDRH3) lengths were identified using IMGT numbering. Somatic hypermutation (SHM) % was calculated from the outputted identity % for each V, D and J region from IMGT, by subtracting the value from 100 %.

To clone R5.034LS, the R5.034 IgG1 heavy chain constant region was replaced with a gene string encoding the same human IgG1 constant heavy region containing the two LS mutations (M451L/N457S) ^33,47^ via the SalI and HindIII restriction sites. Recombinant R5.034 was then produced as previously described.

#### Expression and Purification of Antibodies

Transfections of HEK293F cells using a 1:1 ratio of HC and light chain (LC) plasmids were set up in 10 mL culture volumes in 50 mL vented cap reaction tubes using Expifectamine^TM^ transfection kit (Life Technologies) as per the manufacturer’s instructions. IgG purification was carried out using Econo-PAC® chromatography columns (Bio-Rad) and Protein A Sepharose (Sigma-Aldrich), and purified mAbs were buffer-exchanged into phosphate buffered saline (PBS).

#### Expression and Purification of Proteins

Unless stated otherwise, all PfRH5 and PfCyRPA soluble proteins and reagents were designed based on the 3D7 clone *P. falciparum* reference sequence with all the N-glycan sequons (N-X-S/T) mutated from a serine or threonine to alanine.

##### RH5.1 and RH5ΔNL

The design, production and purification of RH5.1 and RH5ΔNL have previously been described ^16,19,24^. In brief, stable *Drosophila* S2 polyclonal cell lines expressing full-length RH5.1 (residues E26-Q526) or RH5ΔNL (residues K140-K247 and N297-Q526 of PfRH5 with 3D7 or 7G8 *P. falciparum* sequence, differing only by the C203Y polymorphism) were cultured in EX-CELL 420 media and expanded in shake flasks to the desired scale. Cell supernatant was harvested and loaded onto a 10 mL column packed with CaptureSelect affinity C-Tag XL resin, washed in 10 column volumes of TBS pH 7.4 and eluted in 2M MgCl_2_. Fractions were then pooled, concentrated using a 10 kDa Amicon ultra centrifugal filter and run on an HiLoad 16/600 Superdex 200 pg size exclusion chromatography (SEC) column into TBS pH 7.4 (20 mM TRIS-HCl, 150 mM NaCl).

##### RH5ΔNC-Biotin

The production of the RH5ΔNC-Biotin has been described ^38^. Mono-biotinylated RH5ΔNC-Biotin, previously referred to as ‘RH5-Bio’, was generated through co-transfection of Expi293 cells with a plasmid encoding RH5ΔNC-Biotin and another plasmid encoding BirA biotin ligase. RH5ΔNC-Biotin was then purified from the supernatant by C-tag affinity chromatography follow by SEC into TBS pH 7.4 (20 mM TRIS-HCl, 150 mM NaCl).

##### PfCyRPA

Full-length PfCyRPA (residues 29-362) was expressed in Expi293 cells and purified through C-tag affinity chromatography follow by SEC into TBS pH 7.4 as previously described ^25,48^.

##### Basigin

Native human basigin sequence, encoding residues M1-L206, followed by rat CD4 domains 3 and 4 and a C-terminal hexa-histidine tag was expressed through transient transfection of Expi293 cells. Protein was then purified from the supernatant by immobilized metal affinity chromatography (IMAC) using a Ni^2+^ resin followed by SEC into TBS pH 7.4 (20 mM TRIS-HCl, 150 mM NaCl) as previously described ^8,24,49^.

### ELISA

For assessment of antibody binding by ELISA, Nunc Maxisorp plates were coated overnight (>16 h) with either RH5.1, heat-denatured RH5.1 (held at 90 °C for 10 min), or RH5ΔNL at 2 μg/mL. Plates were washed in wash buffer (PBS with 0.05 % Tween 20 [PBST]) and blocked with 100 µL/well of Blocker™ Casein (Thermo Fisher Scientific) for 1 h. Plates were washed and antibodies at 10 µg/mL diluted in casein were added. Following a 2 h incubation, plates were washed and a 1:1000 dilution of goat anti-human IgG (γ-chain specific) alkaline phosphate conjugate antibody (A3187, Thermo Fisher) was added and incubated for 1 h. Plates were washed in washing buffer and 100 µL development buffer (*p*-nitrophenyl phosphate substrate diluted in diethanolamine buffer) was added per well and developed according to internal controls. All mAbs were tested in duplicate against each coating antigen. Unless otherwise stated, 50 µL was added per well and all steps were carried out at RT. A given mAb was tested against all antigens within the same plate.

For the peptide array ELISAs, steps were identical to the above except streptavidin-coated plates (Pierce) were used and were coated with an array of 62 x 20-mer PfRH5 peptides overlapping by 12 amino acids as previously reported ^17^.

#### Antibody Kinetics

High-throughput SPR binding experiments in **Figure 2** and **Figure S2** were performed on a Carterra LSA instrument equipped with a 2D planar carboxymethyldextran surface (CMDP) chip type (Carterra) using a 384-ligand array format. The CMDP chip was first conditioned with 60 s injections of 50 mM NaOH, 1 M NaCl and 10 mM glycine (pH 2.0) before activation with a freshly prepared 1:1:1 mixture of 100 mM MES (pH 5.5), 100 mM sulfo-N-hydroxysuccinimide, and 400 mM 1-ethyl-3-(3-dimethylaminopropyl) carbodiimide hydrochloride. A coupled lawn of goat anti-human IgG Fc (hFc; 50 µg/mL in 10 mM sodium acetate, pH 4.5) (Jackson ImmunoResearch) was then prepared before quenching with 0.5 M ethanolamine (pH 8.5) and washing with 10 mM glycine (pH 2.0). mAbs prepared at 100 ng/mL in Tris-buffered saline with 0.01 % Tween-20 (TBST) were then captured onto the surface in a 384-array format via a multi-channel device, capturing 96 ligands at a time. For binding kinetics and affinity measurements, an eight-point threefold dilution series of RH5.1 protein ending at 100 nM in TBST was sequentially injected onto the chip from lowest to highest concentration. For each concentration, the antigen was injected for 5 min (association phase), followed by TBST injection for 15 min (dissociation phase). Two regeneration cycles of 30 s were performed between each dilution series by injecting 10 mM glycine (pH 2.0) on the chip surface. The SPR results were exported to Kinetics Software (Carterra) and analyzed as nonregenerative kinetics data to calculate association rate constant (*K*_on_), dissociation rate constant (*K*_off_) and equilibrium dissociation constant (K_D_) values via fitting to the Langmuir 1:1 model. Prior to fitting, the data were referenced to the anti-hFc surface then double referenced using the final stabilizing blank injection.

#### Epitope Binning by Surface Plasmon Resonance

High-throughput epitope binning experiments shown in **Figure 1**, **Figure S1** and **Figure S3** were performed in a classical sandwich assay format using the Carterra LSA and an HC30M chip. The chip was conditioned as described above before antibodies prepared at 10 µg/mL in 10 mM sodium acetate (pH 4.5) with 0.05 % Tween were coupled to the surface: the chip surface was first activated with a freshly prepared 1:1:1 activation mix of 100 mM MES (pH 5.5), 100 mM sulfo-N-hydroxysuccinimide, and 400 mM 1-ethyl-3-(3-dimethylaminopropyl) carbodiimide hydrochloride, and antibodies were injected and immobilized onto the chip surface by direct coupling. The chip surface was then quenched with 1 M ethanolamine (pH 8.5), followed by washing with 10 mM glycine (pH 2.0). Sequential injections of 50 nM RH5.1 protein (5 min) followed immediately by the 10 µg/mL sandwiching antibody (5 min), both diluted in HEPES-buffered saline Tween-EDTA (HBSTE) with 0.5 mg/mL bovine serum albumin (BSA), were added to the coupled array and the surfaces regenerated with 10 mM glycine (pH 2.0) using two 30 s regeneration cycles. Data were analyzed using the Carterra Epitope software.

The epitope binning experiment for mouse mAbs QA1 and 9AD4 ^21^ shown in **Figure S4** was performed as above but using a HC200M chip and TBST as the running buffer and diluent.

#### Determination of Protein Blockade by Bio-layer Interferometry (BLI)

All BLI for data shown in **Figure 1** and **Figure S1** was carried out on an OctetRED384 (ForteBio) using anti-human Fc-capture sensors (Sartorius, 18-5060) to immobilize anti-PfRH5 mAbs. Assays were carried out in a 384-well format in black plates (Greiner). For assaying the ability of each mAb to block RH5.1 binding to basigin and PfCyRPA, the experiment followed a sequential assay: mAb immobilization (15 µg/mL, 300 s), RH5.1 binding (1 µM, 300 s), protein ligand binding (3 µM PfCyRPA or basigin, 300 s) with a 30 s dissociation phase in TBST between each step. Finally a 120 s dissociation step was carried out in TBST before a 10 s pulsed regeneration of biosensors with 10 mM glycine (pH 2). Within each plate, PfCyRPA blocking was first assessed for a set of mAbs immediately followed by regeneration and then basigin blocking activity assessed for the same set of mAbs. As internal or “sentinel mAb” controls, each plate included a PfCyRPA-blocking mAb (R5.015), a basigin blocking mAb (R5.004) and a non-blocking mAb (R5.011) ^24,25^. In addition, in each assay, a reference baseline set of biosensors was run in parallel using the same format but replacing the protein ligand binding step with a TBST step.

Data were analyzed in the Octet Data Analysis HT software (Fortebio). The reference biosensors were assigned as references in the software and subtracted from the test biosensors. Steps were aligned to the start of each association step and the association and dissociation was fitted. Response value report points were set at 20 s (start of association) and 290 s (end of association) and exported. For the majority of mAbs, the 290 s report point was used. For a subset of mAbs with very fast dissociation, the 20 s timepoint was used (because report points at 290 s could be erroneously reported as blocking due to dissociation of the underlying RH5.1 surface from the captured mAb). For the RH5.1 report points, the data were unreferenced. For each mAb, the basigin and PfCyRPA responses were normalized by dividing by the RH5.1 response. Any mAb with a normalized response value of <0.04 nm for basigin or PfCyRPA was categorized as “blocking” for that protein ligand. Data were discarded if R5.015 had a response >0.04 nm for PfCyRPA; R5.004 had a response >0.04 nm for basigin; or if R5.011 had a response <0.04 nm for either protein ligand.

#### Assay of Growth Inhibition Activity (GIA)

Single concentration *in vitro* GIA assays were carried as previously described according to the methods of the GIA International Reference Centre at NIAID/NIH, USA ^50^. All assays used 3D7 clone *P. falciparum* parasites cultured in human RBC from in-house volunteer donations or supplied by the UK NHS Blood and Transplant service for non-clinical issue. Briefly, mAbs were buffer exchanged into incomplete parasite growth media (ICM = RPMI, 2 mM L-glutamine, 0.05 g/L hypoxanthine, 5.94 g/L HEPES) before performing the GIA assay and allowing parasites to go through a single cycle of growth. To ensure consistency between experiments, in each case the activity of a negative control human mAb, EBL040 ^51^ which binds to the Ebola virus glycoprotein, and three anti-PfRH5 mAbs with well-characterized GIA (2AC7, QA5, and 9AD4; or 2AC7, R5.016, and R5.034 ^21,24^) were run alongside the test mAbs, and were all tested in triplicates. mAbs showing >30 % GIA were subsequently tested in an eight step, five-fold dilution series with a final assay start concentration of 2 mg/mL to determine interpolated EC values. The resultant data were transformed according to x=log(x) and the transformed data were fitted by four-parameter non-linear regression. GIA values were interpolated from the resultant curve. If a mAb did not reach a sufficiently high GIA (i.e. the mAb did not reach 30 %, 50 % or 80 % at any test concentration), then it was assigned a “negative” value of 10,000 µg/mL for that particular EC readout.

For screening of intra-PfRH5 mAb interactions, shown in **Figure 5A and Figure S5A**, single concentration assays were carried out as above, with neutralizing antibodies added at a concentration equivalent to their interpolated EC_50_ value and non-neutralizing antibodies held at 0.3 mg/mL. For testing of non-neutralizing antibodies in combination, shown in **Figure 5B**, both mAbs in a pair were held at 0.2 mg/mL. For dilution curves, titrated mAbs were set up in a 7-step, five-fold dilution series starting at 0.5 mg/mL per mAb. To each titrated mAb dilution, a held concentration of a second mAb (or a premixed combination of two mAbs) was added at a final concentration of 0.2 mg/mL per held mAb. For each curve, a well containing the second mAb (or premixed combination) at the test concentration was set up alone within the same assay plate. Curve fitting and data processing was carried out as above.

For mixed titration curves, shown in **Figure 5C** and **Figure S5E-G**, two or more mAbs were premixed and set up in an 8-step, five-fold dilution series starting at 0.5 mg/mL per test mAb. Curve fitting and data processing was carried out as above.

For synergy curves with polyclonal antibody shown in **Figure S5D**, total IgG (purified from the serum of VAC063 vaccinees ^13^) was set up as above at a starting concentration of 14 mg/mL total IgG in a 7-step 2-fold dilution curve.

For analysis of synergistic or antagonistic interactions, the Bliss additivity ^52^ was determined based on the measured activity from each antibody alone (1 and 2) using the following formula:

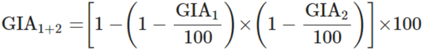

#### Purification of IgG

For the polyclonal antibody (pAb) pool used in **Figure S5D**, a pool of human serum from the VAC063 clinical trial ^13^ was filtered through a 0.22 µM syringe filter (Milipore) and diluted 1:1 in PBS. Total IgG was then purified using a HiTrap Protein G column (Cytiva) on an ÄKTA Pure chromatography system (Cytiva), and the eluted total IgG was buffer exchanged into ICM using a centrifugal concentrator with a 30K MWCO (Cytiva). Total IgG concentration was determined by reading absorbance at 280 nm on a Thermo Scientific™ NanoDrop™ OneC Spectrophotometer. IgG was then depleted for anti-RBC antibodies by the addition of 1 µg packed 100 % hematocrit RBC per 1 μg IgG and incubated at RT with agitation for 1 h. RBC were then pelleted by centrifugation at 1,000 x*g* for 5 min and the supernatant removed. Anti-RH5.1 antibody titers or concentrations were determined by standardized quantitative ELISA methodology as previously reported ^17^.

#### Antibody Sequence Annotation

All annotation of antibody heavy and light chain gene sequences was performed using IMGT V-Quest program version 3.5.31 using default parameters. The IGH locus was selected for heavy chain sequences, IGL for λ light chains and IGK for κ light chains. F+ORF+ in-frame P was used as the IMGT/V-QUEST reference directory set and the option to search for insertions and deletions was selected.

#### Design of Public Clonotype Germline Revertant mAbs

Synthetic antibodies shown in **Figure 3E** were designed based on the HV3-7/LV1-36 gene combination. For the germline heavy chains (“HC GL”), amino acids E1 to R98 of the V-region of each public clonotype mAb (R5.034, R5.102, R5.237 and R5.270) were replaced with amino acids E1 to R98 of the germline HV3-7*01 F gene (IMGT accession number: M99649) V-region obtained from IMGT. The sequence for each mAb from R98 was unchanged from the wildtype (WT) mAb, resulting in four germline HV3-7 heavy chains, each biased with the CDRH3 J- and D-regions of the respective public clonotype mAb.

For the germline LV1-36 light chain (“LC GL”), amino acids Q1 to G98 of the germline LV1-36*01 F gene (IMGT accession number Z73653) V-region were concatenated with amino acids VVFGGGTKLTVL of the germline LJ2*01 F gene (IMGT accension number M15641). Note that R5.034, R5.102 and R5.270 were equally likely to use the LJ2*01 F or LJ3*01 F (IMGT accession number M15642) gene segments, only R5.237 had a greater identity to the LJ3*02 (IMGT accession number D87023) gene segment.

Synthetic genes were cloned into AbVec expression vectors as described above. For the production of heavy chain revertant antibodies (HC GL), HEK293F cells were transfected with expression plasmids containing one of the four germline HV3-7 heavy chains, and the cognate WT light chain of the respective public clonotype mAb. For the production of light chain revertant antibodies (LC GL), cells were transfected with expression plasmids containing one of the four WT public clonotype mAb heavy chains and their respective germline LV1-36 light chain. For the production of full revertants (“Full GL”), cells were transfected with expression plasmids containing one of the four germline HV3-7 heavy chains and the respective germline LV1-36 light chain.

To generate the public clonotype CDRH3 knockout antibodies (“CDRH3_KO”), the 9 amino acids from positions 99 to 107 in the CDRH3 of each antibody (IMGT numbering) were replaced with a ‘randomized’ sequence of amino acids with a matched length – HQSGKLVNMN. No amino acid in this sequence was conserved with respect to the equivalent position in any of the public clonotype CDRH3 sequences. The ‘randomized’ region was chosen to test the effects of altering amino acids derived from somatic hypermutation of rearrangement, whilst preserving the largely conserved and germline templated sequences at the N-(positions 97-98) and C-(positions 108-109) termini of the 13 amino acid CDRH3. CDRH3 knockout heavy chains were cloned into expression vectors as described above. Cells were transfected with expression plasmids containing the respective CDRH3 knockout heavy chain for each public clonotype antibody and its cognate WT light chain.

#### Fab Cloning and Expression

To express recombinant Fabs, heavy chain plasmids were generated using primers to amplify the heavy chain VH and CH1 domain only (5’ – GAG GAT GGT CAT GTA TCA TC and 5’ – CGC AAG CTT CTA AGT TTT GTC ACA AGA TTT GGG C) and then used to transfect Expi293F cells in combination with the corresponding light chain plasmids. Fab-containing supernatants were purified by affinity chromatography with either a HiTrap LambdaFabSelect column (17548211, Cytiva) or a HiTrap Protein G column (17040501, Cytiva) on an ÄKTA Pure chromatography system as per manufacturer’s instructions.

#### Production of RH5ΔNL:Fab Complexes for X-ray Crystallography

Fabs and RH5ΔNL protein (7G8 sequence with Y203) were produced as described above, and buffer exchanged into HEPES-buffered saline (HBS, 25 mM HEPES, 150 mM NaCl, pH 7.5) using a 10K MWCO centrifugal concentrator device (Cytiva). RH5ΔNL was subjected to limited proteolysis at a concentration of 1-2 mg/mL, with the addition of 1 µL GluC protease (P8100S, NEB) at 1 mg/mL per mg of RH5ΔNL (1:1000 wt/wt). Protein was incubated at RT for 4-16 h. Fabs were then added at a 5 % molar excess. The complex was incubated for 1 h at RT. The complex was then subjected to surface lysine methylation by the addition of 20 µL per mL of complex of 1 M dimethylamine borane complex (ABC) (Santa Cruz Biotechnology, sc-252506) and 40 µL per mL of complex of 1 M formaldehyde prepared from 37 % stock (Thermo Fisher, 10630813). The complex was incubated for 1 h at RT, followed by a further addition of 20 µL ABC and 20 µL formaldehyde per mL of complex. After a further 1 h incubation, 20 µL ABC per mL of complex was added and the solution was incubated at 4 °C overnight. Aggregates were removed by spinning the sample in a centrifugal filter device (UFC40GV00, Milipore) at 4000 x*g* for 10 min. The flow-through was concentrated to <2 mL if necessary and run onto a Superdex 16/600 200 pg SEC column (Cytiva) that had been pre-equilibrated in Tris-buffered saline (25 mM Trizma, 150 mM NaCl, pH 7.4). Complex containing fractions were pooled and concentrated to 8-12 mg/mL using a centrifrugal concentrator with a 10K MWCO (Cytiva) for crystal screening.

#### Structure Determination by X-ray Crystallography

All crystallization was conducted using vapour diffusion in MRC 3 Lens sitting drop crystallization plates (SwisSci, High Wycombe, UK) and 150 nL drops (ratio 100 nL protein solution (8 mg/µL:50 nL screen condition) dispensed using a Mosquito nano-pipetting robot (STP Labtech, Melbourn, UK). Crystallization plates were incubated at 20 °C with crystals appearing between 4 and 28 days. The crystals were mounted with LithoLoops (Molecular Dimension, Rotherham, UK) using the CrystalShifter crystal harvesting robot ^53^ (Oxford LabTech, UK) and cryo-protected in a solution of 25% Ethylene Glycol.

Diffraction quality crystals of complex of RH5ΔNL:R5.034:R5.028 Fabs were obtained from PEG/Ion Screen (Hampton Research, Aliso Viojo, CA) condition H6 (0.02 M Calcium chloride dihydrate, 0.02 M Cadmium chloride hydrate, 0.02 M Cobalt(II) chloride hexahydrate, 20 % w/v Polyethylene glycol 3,350). All diffraction data were collected at Diamond Light Source (Proposal ID: mx28172). Initially samples were sent to i03 for Unattended Data Collection (Native Experiment; 12.7 keV, 1.7 Å, 2 x 360° sweeps, 1st at chi=0 and 2nd at chi=30), before being transferred to i24 for manual data collection (4.00Å, 360°).

Diffraction quality crystals of complex RH5ΔNL:R5.251 Fab were obtained from the PurePEGs (Anatrace, Maumee, OH) condition B8 (0.3M Calcium chloride, and 0.1M Magnesium formate HCl 6, 22.5% (v/v) PurePEGs Cocktail). Data were collected at i03 (Diamond Light Source (Proposal ID: mx28172)) using Unattended Data Collection (Native Experiment; 12.7 keV, 1.7 Å, 2 x 360° sweeps, 1st at chi=0 and 2nd at chi=30).

#### Data Processing and Model Refinement

Datasets for RH5ΔNL:R5.251 and RH5ΔNL:R5.034:R5.028 were manually processed on the Diamond Cluster using DIALSs ^54^, confirming the auto-processed xia2.DIALS ^55^ output from SynchWeb/iSpyB ^56,57^. All subsequent processing was done on local computers using the CCP4i GUI ^58^ or on CCP4 Cloud ^59^. RH5ΔNL:R5.251 and RH5ΔNL:R5.034 data were truncated at 3.2 Å and 4 Å, respectively. Although the apparent crystal symmetry was orthorhombic, refinement failed to reduce R-free as would usually be expected. Data for the putative complex of RH5ΔNL:R5.034:R5.028 were reprocessed in P21 with a beta angle of ∼90° and pseudomerohedral twinning is suspected; however, there was no evidence of R5.028 Fab in the data, suggesting it was lost during crystal formation with just RH5ΔNL:R5.034 remaining. Molecular replacement was done with Phaser ^60^ using homology model coordinates downloaded from the Protein Data Bank (PDB) based on a sequence similarity search ^61^. The following homology models were used for: i) RH5ΔNL (PDB: 4WAT); and ii) R5.251 Variable, Heavy (PDB: 6OC7); Constant, Heavy (PDB: 6OC7); Variable, Light (PDB: 5XKU); Constant, Light (PDB: 5XKU); and iii) R5.034 Variable, Heavy (PDB: 5X8M); Constant, Heavy (PDB: 1BJ1); Variable, Light (PDB: 5WL2); Constant, Light (PDB: 3H0T). Subsequent model refinement and building was performed in Coot ^62^ and REFMAC ^63^. Once stable models had been built they were put through the PDB-REDO ^64^ pipeline to be optimized, and LORESTR ^65^ for further low-resolution refinement.

#### Computational Prediction of GIA from mAb Gene Usage

Subject gene usage was one-hot encoded using the R package mltools (v0.3.5). To more robustly assess the relationship between gene usage and GIA %, only pairs of genes that occurred in at least 4 individuals were considered. Further filtering was applied to select only gene pairs whose mean associated GIA % was outside a conservatively selected 33 % deviation from the dataset mean (below 42 % GIA %, or above 82 % GIA %). An additional requirement was added that GIA % values associated with gene pairs must be either entirely above or below the dataset mean and not spanning. To assess statistical significance, 1000 rounds of permutation were run, where gene use was scrambled across the dataset and the same filtering was performed to search for significant gene pairs. A *P* value (*P* < 0.001) was calculated from the tail probability of the generated null distribution. An R markdown for the analysis is available.

#### R5.034 and R5.034LS Binding Kinetics to RH5.1

For **Figure S6A**, SPR was carried out using the Biacore™ X100 machine and software. Purified recombinant R5.034 or R5.034LS was immobilized on Sensor Chip Protein G (Cytiva) through a 30 s injection of 16 nM antibody. RH5.1 protein was diluted in PBS + P20 running buffer (137 mM NaCl, 2.7 mM KCl, 10mM Na_2_HPO_4_, 1.8 mM KH_2_PO_4_, 0.005 % surfactant P20 (Cytiva)) to yield a final concentration of 15.6 nM. Samples were injected for 180 s at 30 μL/min before dissociation for 800 s. The chip was then regenerated with a 45 s injection of 10 mM glycine pH 1.5. Antibody kinetics were determined through a two-fold, five-step dilution curve. Data were analyzed using the Biacore X100 Evaluation software v2.0.2. A global Langmuir 1:1 interaction model was used to determine antibody kinetics.

#### R5.034 and R5.034LS Binding Kinetics to Human FcRn

For **Figure 6C**, SPR was carried out using the Biacore™ X100 machine and software. Recombinant human FcRn protein (Acro Biosystems) was immobilized onto a CM5 Sensor Chip (Cytiva) using the standard amine coupling protocol yielding ∼200 response units (RU). R5.034 and R5.034LS were diluted to a final concentration of 6.4 µM and 1.6 µM, respectively, in either MES pH 6.0 (20 mM MES, 150 mM NaCl) or TBS pH 7.4 (20 mM TRIS HCl, 150 mM NaCl) running buffer. Samples were injected for 180 s at 30 μL/min before dissociation for 600 s. The chip was regenerated with a 30 s injection of PBS pH 7.4. Affinity was determined using a two-fold, nine-step dilution curve. Data were analyzed using the Biacore X100 Evaluation software v2.0.2, and the equilibrium dissociation constant was determined from a plot of steady state binding levels.

#### *P. falciparum* Sporozoite Challenge in Liver-Chimeric Humanized Mice

The FRG huHep mouse studies in **Figure 6E** were conducted similar to studies previously published ^66^ with modifications. FRG huHep mice on the NOD background were purchased from Yecuris, Inc. (Beaverton, OR, USA). Mice were pre-screened to have a serum human albumin level indicative of >90 % humanization of hepatocytes. Mice were then infected with *P. falciparum* NF54 strain via mosquito bite. Mosquitos were purchased from the Johns Hopkins Malaria Research Institute Insectary Core and used only if >50 % of mosquitos were infected with a mean of >10 oocysts per midgut. Based on this midgut prevalence and/or salivary gland “smash test” (dissection of individual mosquito salivary glands followed by microscopic observation of sporozoites), mosquitos were apportioned to cages equivalent to 5 infectious mosquitos per mouse. Mosquitos were then allowed to feed on mice anesthetized under isoflurane for 10 min, with lifting of mice every minute to encourage probing as opposed to blood feeding.

On day 5 post-infection, mice were intravenously injected with both mAb and human RBC. Monoclonal antibodies (either 625 µg anti-PfRH5 human IgG1 mAb R5.034 or 675 µg human IgG1 negative control mAb 1245 against the sexual-stage malaria antigen Pfs25 ^67^) were delivered via the retro-orbital route diluted to 100 μL total volume in sterile PBS. Human RBC were obtained from a commercial vendor (BloodWorks Northwest, Seattle, WA, USA) and washed three times with sterile RPMI to remove serum and white blood cells. Human RBC were injected via the tail vein in a total volume of 400 μL containing 50 μL clodronate liposomes (Formumax Cat #F70101C-AH), 5μL penicillin/streptomycin and 345 μL 70 % hematocrit human RBC (packed RBC diluted to 70 % hematocrit with sterile RPMI).

On day 6 post-infection, mice were bled via the retro-orbital plexus using non-heparinized capillary tubes. Blood was transferred to 1.5 mL Eppendorf tubes and allowed to clot for 30 min at room temperature (RT). Serum was separated by centrifugation in a table top centrifuge at 9600 rpm for 10 min and stored at -80 °C until use. Mice were then injected with 700 μL 70 % hematocrit human RBC. Injection of this volume of human RBC was repeated on days 9 and 11.

On days 7, 9, 11 and 13 post-infection, mice were bled via the retro-orbital plexus using heparinized capillary tubes and 100 μL whole blood was transferred to 1.9 mL nucliSENS Lysis Buffer (Biomerieux Inc. Cat# 200292). Blood was allowed to lyse at RT for at least 30 min prior to storage at -80 °C before qRT-PCR analysis. Terminal serum was also collected on day 13 via cardiac puncture into a 1 mL syringe with no anticoagulant with separation performed as above. Blood samples were then blinded and sent to the lab of Dr. Sean Murphy at the University of Washington for quantification of *Plasmodium* 18s rRNA following published methods ^68^.

Serum levels of R5.034 antibody were determined by testing dilutions of test sera on a RH5.1 protein capture ELISA including use of a standard curve of purified recombinant R5.034 mAb. In brief, ELISA plates were coated with RH5.1 protein at 2 µg/mL and then blocked with Blocker™ Casein solution. Test sera were diluted and then titrated using a 12-point dilution curve with 1:2 dilutions, and read off an R5.034 standard curve to determine concentration.

### STATISTICAL ANALYSIS

Analysis was performed using GraphPad Prism version 10.0.2 (GraphPad Software, LLC). Tests and statistics are described in Figure Legends. Non-parametric tests were chosen for non-normally distributed data. To determine EC values, mAb dilution curves were transformed according to x=log(x) and the transformed data were fitted to a curve by four-parameter non-linear regression. GIA values were interpolated from the resultant curve with upper and lower 95 % confidence intervals. If a mAb did not reach a sufficiently high GIA (i.e. the mAb did not reach 30 %, 50 % or 80 % GIA at any test concentration), then it was assigned a “GIA-negative” value of 10,000 µg/mL for the purposes of data visualization and statistical testing. In all statistical tests, reported *P* values are two-tailed and *P* < 0.05 considered significant.

