## Supplementary Material for "Analysis of the Diverse Antigenic Landscape of the Malaria Invasion Protein RH5 Identifies a Potent Vaccine-Induced Human Public Antibody Clonotype"

### Supplementary Material for Barrett JR et al.

### Figure S1.

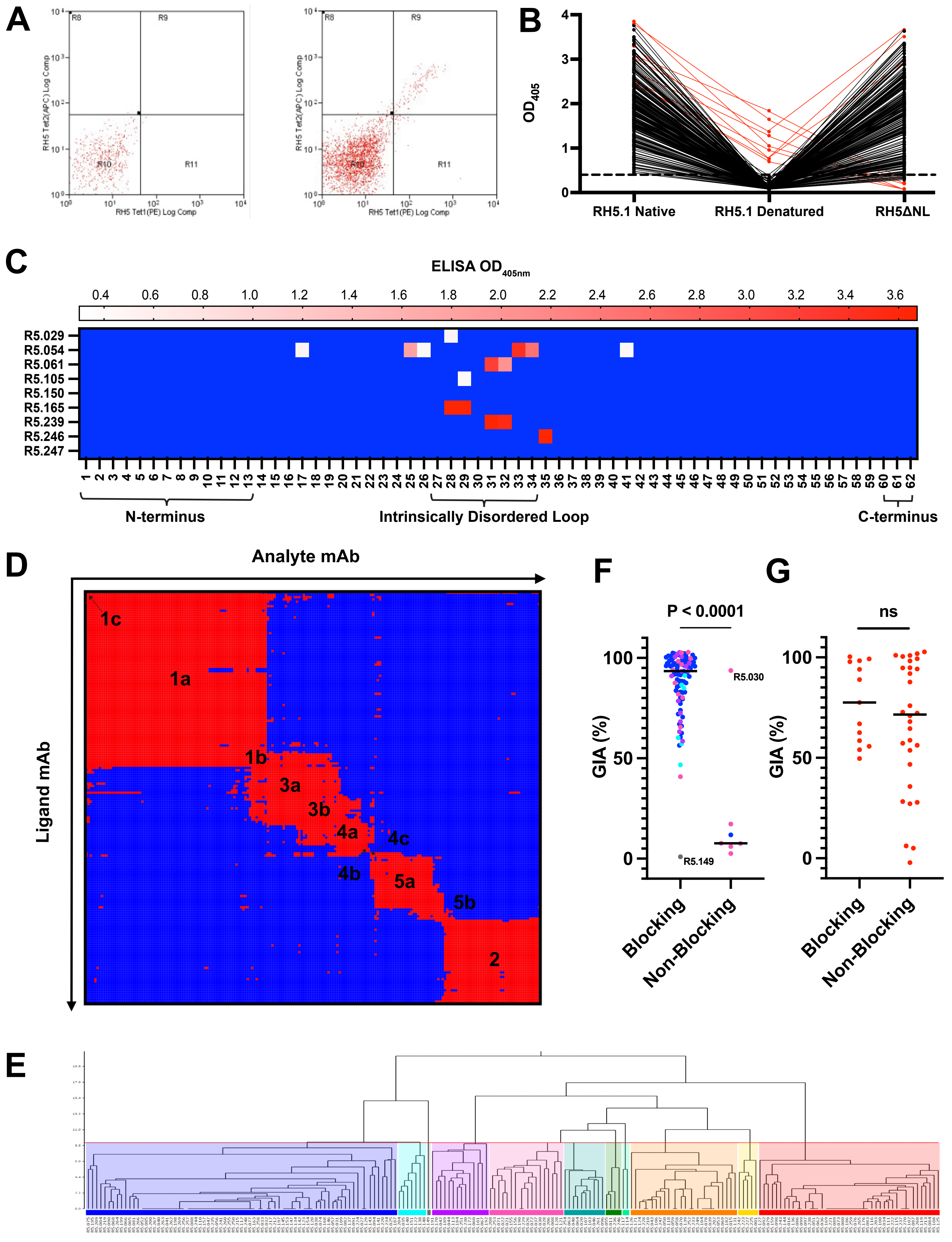

A
B
RH5 Tet2(APC) Log Comp
RH5 Tet1(PE) Log Comp
R8
R9
R10
R11
OD405
RH5.1 Native
RH5.1 Denatured
RH5ΔNL
C
ELISA OD405nm
0.4 0.6 0.8 1.0 1.2 1.4 1.6 1.8 2.0 2.2 2.4 2.6 2.8 3.0 3.2 3.4 3.6
R5.029
R5.054
R5.061
R5.105
R5.150
R5.165
R5.239
R5.246
R5.247
N-terminus
Intrinsically Disordered Loop
C-terminus
D
Analyte mAb
Ligand mAb
1c
1a
1b
3a
3b
4a
4c
4b
5a
5b
2
F
P < 0.0001
GIA (%)
R5.030
R5.149
Blocking
Non-Blocking
G
ns
GIA (%)
Blocking
Non-Blocking
E

### Figure S1: Isolation and epitope mapping of anti-PfRH5 human mAbs; related to Figure 1.

(**A**) Example gating strategy for the sorting of PfRH5-specific B cells from within the live, single, CD19+ IgG+ lymphocyte population. Lymphocytes co-staining with monobiotinylated-RH5ΔNC conjugated to streptavidin-PE (PfRH5-PE probe = RH5 Tet1(PE)) and monobiotinylated-RH5ΔNC conjugated to streptavidin-APC (PfRH5-APC probe = RH5 Tet2(APC)) in quadrant R9 were sorted for mAb production. For each sorting experiment, a negative control sample (i.e. PBMC from a PfRH5-naïve volunteer) was used to set the quadrant gate (left hand plot) before sorting the PfRH5 vaccinee (right hand plot). (**B**) Results of ELISA on mAbs using Native RH5.1, heat-denatured RH5.1, or native RH5ΔNL proteins. Lines between dots connect results for individual mAbs. A dotted line is shown at the positive cut-off of 0.3 $OD_{405}$. Red lines indicate the mAbs that recognized denatured RH5.1. (**C**) Results of linear epitope mapping by ELISA using an array of 62 20-mer overlapping peptides as previously reported [17]. Each mAb was run in singlicate against each peptide. The mean background of N=4 PBS-coated control wells was subtracted from the response for each mAb. Negative values below the cut-off of 0.3 $OD_{405nm}$ are shown in blue. Peptides are annotated with the linear region of PfRH5 to which they correspond. (**D**) Heat map displaying competition interactions between mAbs as ligands (y-axis) and analytes (x-axis). Red squares indicate competition between the mAb pair, blue squares indicate no-competition between the two mAbs. (**E**) Competition profiles were used to cluster mAbs into epitope supercommunities and communities represented by colored regions of the dendrogram (color-coded as per **Figure 1A**) using the McQuitty clustering method in the Carterra Epitope software. (**F**) The GIA of mAbs against 3D7 clone *P. falciparum* (tested at high concentration of 0.8-2 mg/mL) from supercommunities 1 and 3 combined (N=113), or (**G**) community 2 (N=43), categorized by each mAb's ability to block basigin as measured by BLI. Data show individual mAbs and the median. Significance tested by Mann-Whitney test.

### Figure S2.

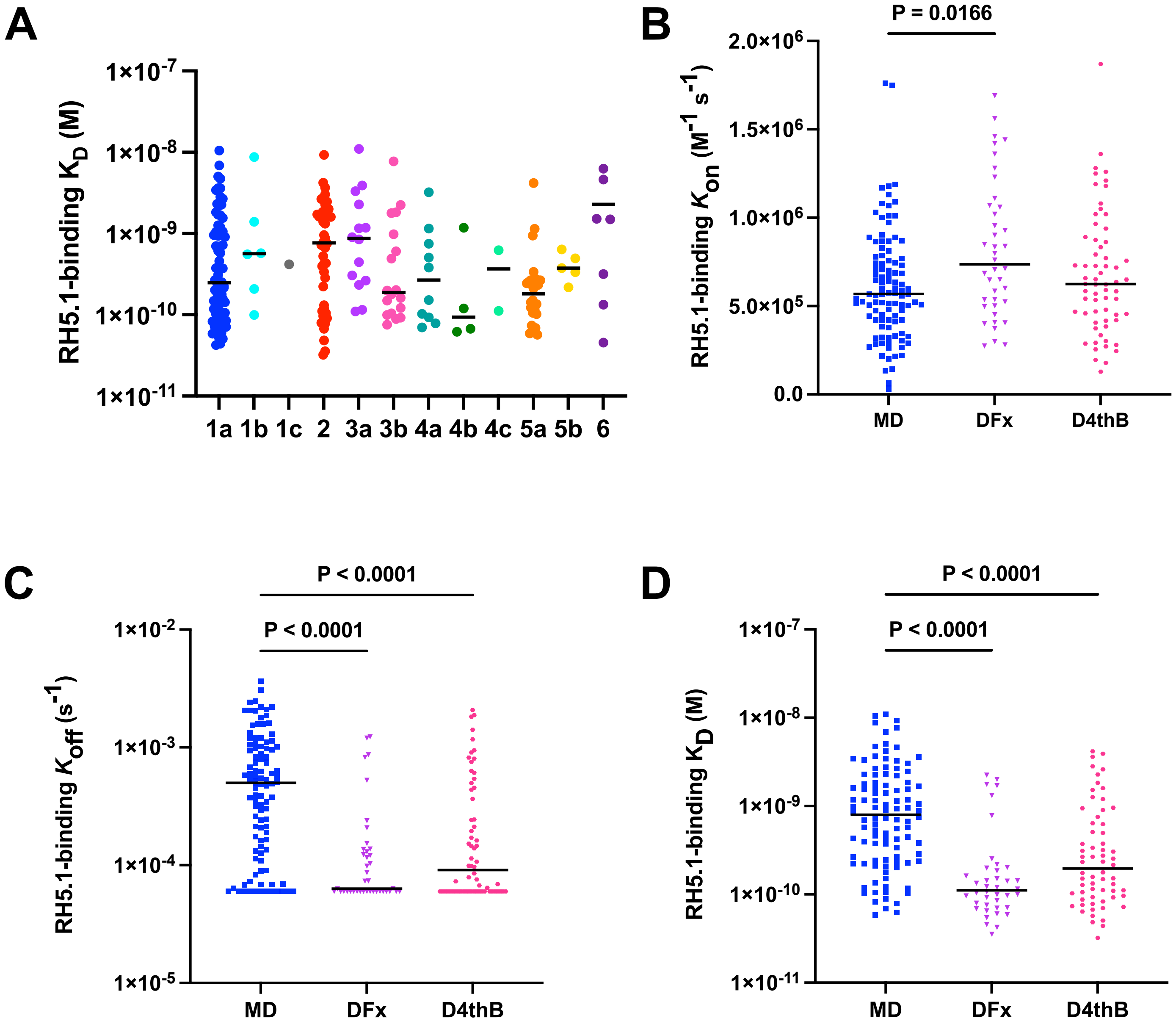

A
RH5.1-binding K_D (M)
1×10-7
1×10-8
1×10-9
1×10-10
1×10-11
1a 1b 1c 2 3a 3b 4a 4b 4c 5a 5b 6
B
P = 0.0166
RH5.1-binding K_on (M-1 s-1)
2.0×10^6
1.5×10^6
1.0×10^6
5.0×10^5
0.0
MD DFx D4thB
C
P < 0.0001
P < 0.0001
RH5.1-binding K_off (s-1)
1×10-2
1×10-3
1×10-4
1×10-5
MD DFx D4thB
D
P < 0.0001
P < 0.0001
RH5.1-binding K_D (M)
1×10-7
1×10-8
1×10-9
1×10-10
1×10-11
MD DFx D4thB

### Figure S2: Binding affinities of anti-PfRH5 mAb epitope communities; related to Figure 2.

(**A**) The binding affinity ($K_D$) to RH5.1 (full-length PfRH5 protein) was determined by HT-SPR. Results from **Figure 2A** are shown plotted by epitope community as defined in **Figure 1A**. Individual mAbs are shown and the line is shown at the median for each community. (**B**) Association rate ($K_{on}$), (**C**) dissociation rate ($K_{off}$) and (**D**) binding affinity ($K_D$) constants of mAbs were plotted by the RH5.1/AS01$_B$ dosing regimen of the vaccinated volunteer from which they were derived in the Phase 1/2a clinical trial [1]. The mAbs were isolated from vaccinees after their final immunization, either given in a monthly dosing (MD) schedule (N=102), or following a delayed fractional booster dose (DFx ; N=40) or a delayed fourth boost (D4thB ; N=64) with a 4-5 month interval between the prior vaccination and the final vaccination, as previously reported [1]. *P* values show significant differences between groups as determined by a Kruskal-Wallis test with Dunn's multiple comparison post-test. Lines are shown at the median.

### Figure S3.

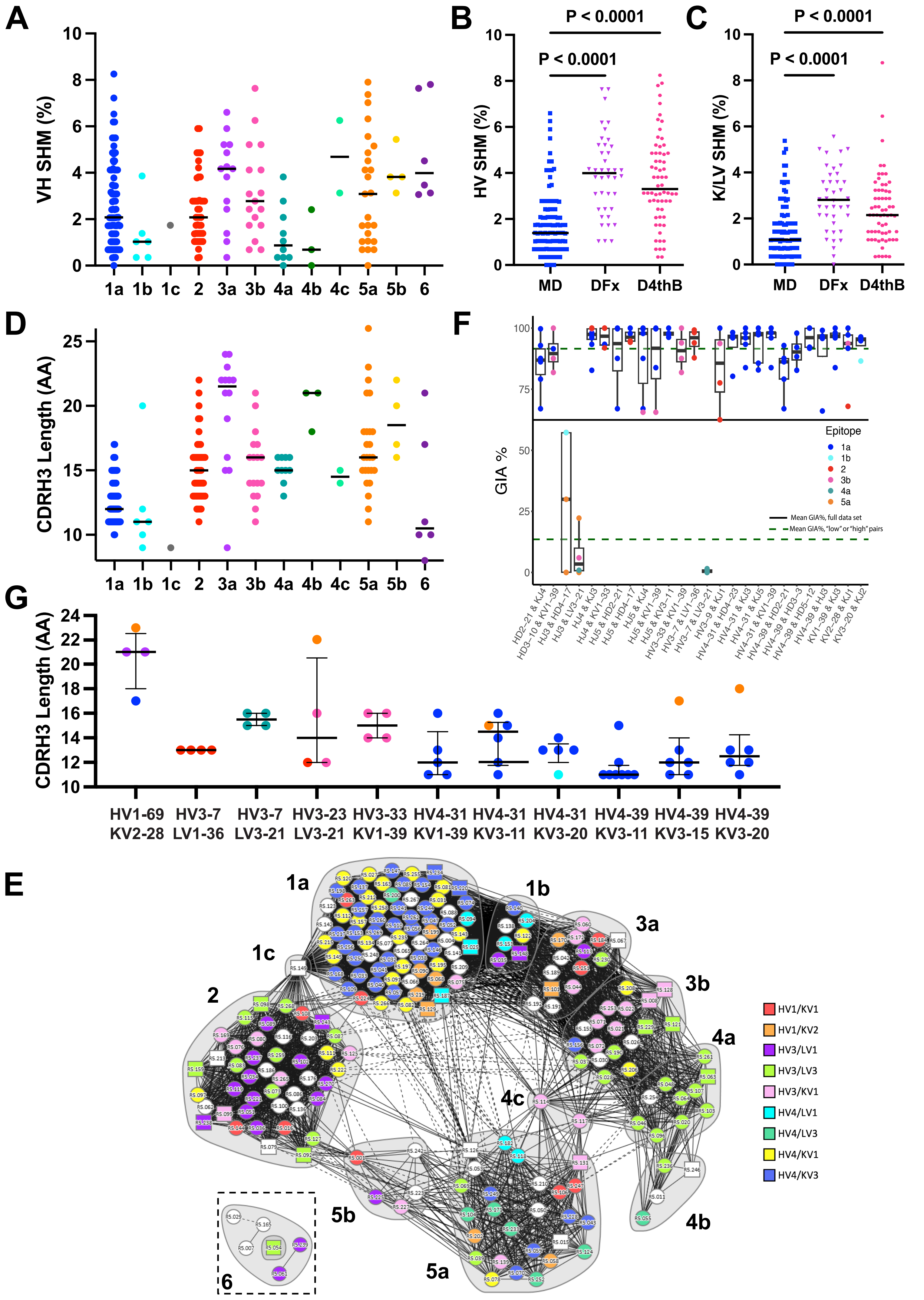
A
VH SHM (%)
1a 1b 1c 2 3a 3b 4a 4b 4c 5a 5b 6
B
P < 0.0001
P < 0.0001
HV SHM (%)
MD DFx D4thB
C
P < 0.0001
P < 0.0001
K/LV SHM (%)
MD DFx D4thB
D
CDRH3 Length (AA)
1a 1b 1c 2 3a 3b 4a 4b 4c 5a 5b 6
F
GIA %
Epitope
1a
1b
2
3b
4a
5a
Mean GIA%, full data set
Mean GIA%, "low" or "high" pairs
G
CDRH3 Length (AA)
HV1-69 KV2-28
HV3-7 LV1-36
HV3-7 LV3-21
HV3-23 LV3-21
HV3-33 KV1-39
HV4-31 KV1-39
HV4-31 KV3-11
HV4-31 KV3-20
HV4-39 KV3-11
HV4-39 KV3-15
HV4-39 KV3-20
E
1a
1b
1c
2
3a
3b
4a
4b
4c
5a
5b
6
HV1/KV1
HV1/KV2
HV3/LV1
HV3/LV3
HV3/KV1
HV4/LV1
HV4/LV3
HV4/KV1
HV4/KV3

### Figure S3: Sequence analysis of anti-PfRH5 mAbs; related to Figure 3.

(**A**) Percentage SHM in the HV gene segment are shown plotted by epitope community as defined in **Figure 1A**. Individual mAbs are shown and the line at the median for each community. (**B**) Percentage SHM in the HV heavy or (**C**) KLV light chain gene segments were plotted by the RH5.1/AS01$_B$ dosing regimen of the vaccinated volunteer they were derived from in the Phase 1/2a clinical trial [1]. The mAbs were isolated from vaccinees after their final immunization, either given in a monthly dosing (MD) schedule (N=102), or following a delayed fractional booster dose (DFx ; N=40) or a delayed fourth boost (D4thB ; N=64) with a 4-5 month interval between the prior vaccination and the final vaccination, as previously reported [1]. *P* values show significant differences between groups as determined by a Kruskal-Wallis test with Dunn's multiple comparison post-test. Lines are shown at the median. (**D**) As for panel **A**, except data show the CDRH3 length in number of amino acids (AA). (**E**) Community network plot overlaid with anti-PfRH5 mAb heavy and light chain gene family pairings. The most common pairings are indicated by the color legend, mAbs with pairings used by fewer than 5 mAbs are colored white. (**F**) An unbiased computational modelling analysis of all available antibody gene sequence data across the anti-PfRH5 mAb panel was used to identify combinations of heavy and light chain variable gene segment usage that are predictive of high GIA. A black line is shown at the mean GIA % of the full dataset. Green dotted lines are shown at the mean GIA % of mAbs classed as "high" (above the mean GIA % of the full dataset) or low (below the mean GIA % of the full dataset). Antibody epitope community is colored as defined in **Figure 1A**. (**G**) Gene pairs in the anti-PfRH5 panel with N≥4 representative mAbs plotted in groups along with their CDRH3 length in number of amino acids (AA). Each mAb is colored by its epitope community as defined in **Figure 1A**. Lines show the median and error bars show the inter-quartile range.

### Figure S4.

A

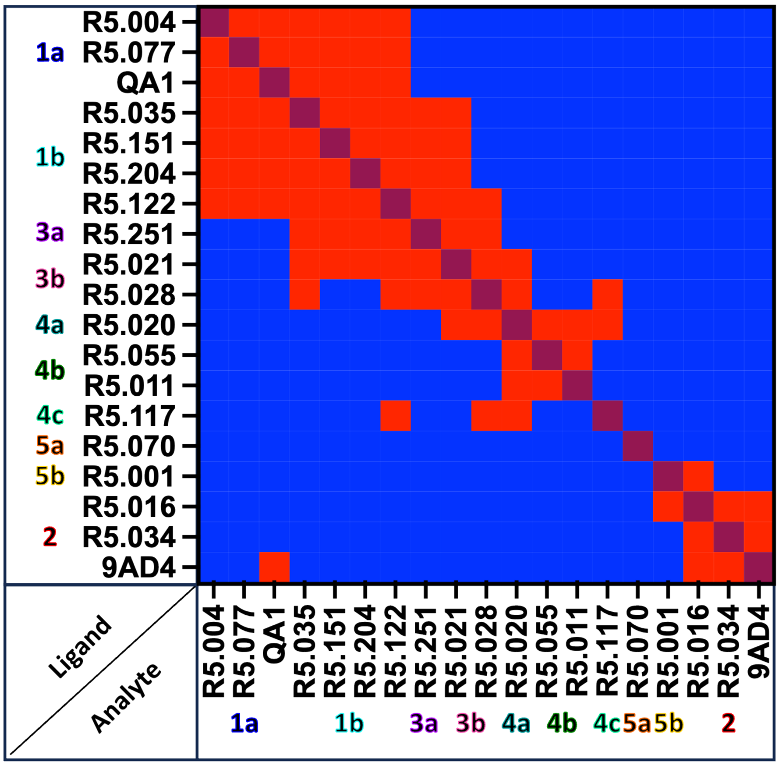

R5.004
1a R5.077
QA1
R5.035
1b R5.151
R5.204
R5.122
3a R5.251
3b R5.021
R5.028
4a R5.020
4b R5.055
R5.011
4c R5.117
5a R5.070
5b R5.001
R5.016
2 R5.034
9AD4
Ligand
Analyte
R5.004 R5.077 QA1 R5.035 R5.151 R5.204 R5.122 R5.251 R5.021 R5.028 R5.020 R5.055 R5.011 R5.117 R5.070 R5.001 R5.016 R5.034 9AD4
1a 1b 3a 3b 4a 4b 4c 5a 5b 2


B

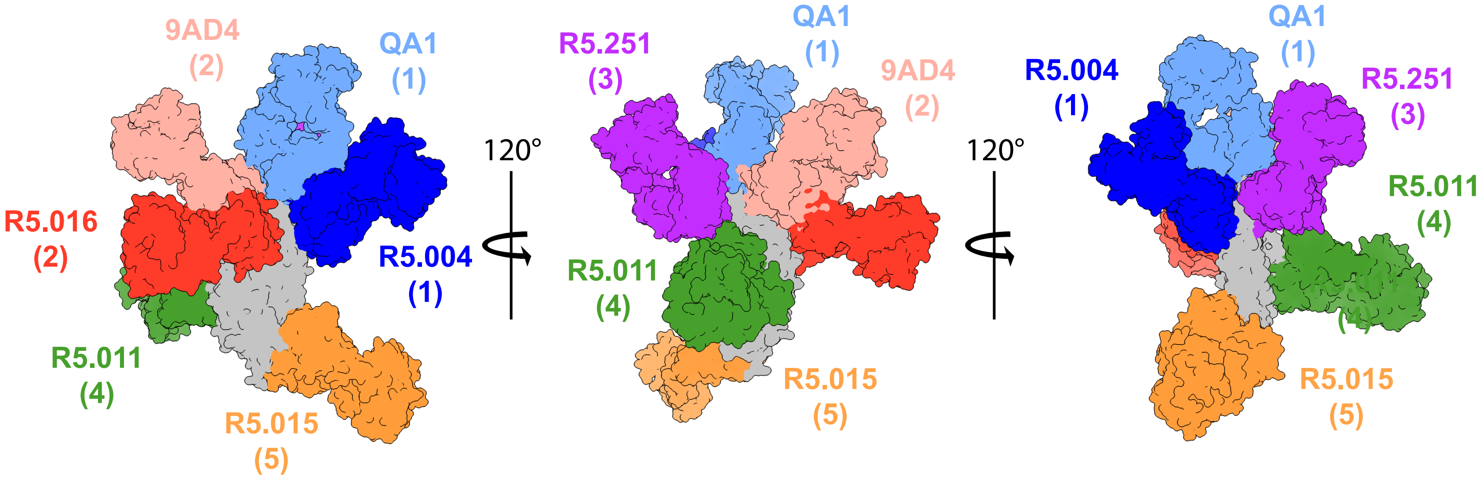

9AD4 (2)
QA1 (1)
R5.016 (2)
R5.004 (1)
R5.011 (4)
R5.015 (5)
120°
R5.251 (3)
QA1 (1)
9AD4 (2)
R5.011 (4)
R5.015 (5)
120°
R5.004 (1)
QA1 (1)
R5.251 (3)
R5.011 (4)
R5.015 (5)

### Figure S4: Epitope mapping murine mAbs and structures of all anti-PfRH5 mAbs; related to Figure 4.

(**A**) Heat map displaying competition interactions between mAbs as ligands (y-axis) and analytes (x-axis) as determined by HT-SPR and including the murine mAbs QA1 and 9AD4 [2]. Red squares indicate competition between the mAb pair, blue squares indicate no-competition between the two mAbs and maroon squares denote self-competition. (**B**) Crystal structure of PfRH5 using RH5ΔNL protein (grey) bound to R5.251 Fab fragment (violet, community 3a), aligned with the structures of Fabs R5.004 (blue, community 1a, PDB ID: 6RCU); R5.016 (red, community 2, PDB ID: 6RCU) and R5.011 (green, community 4b, PDB ID: 6RCV) [3]; R5.015 (orange, community 5b, PDB ID: 7PHU) [4]; and QA1 (pale blue, community 1a, PDB ID: 4U1G) and 9AD4 (pale red, community 2, PDB ID: 4U0R) [5].

**Figure S5.**

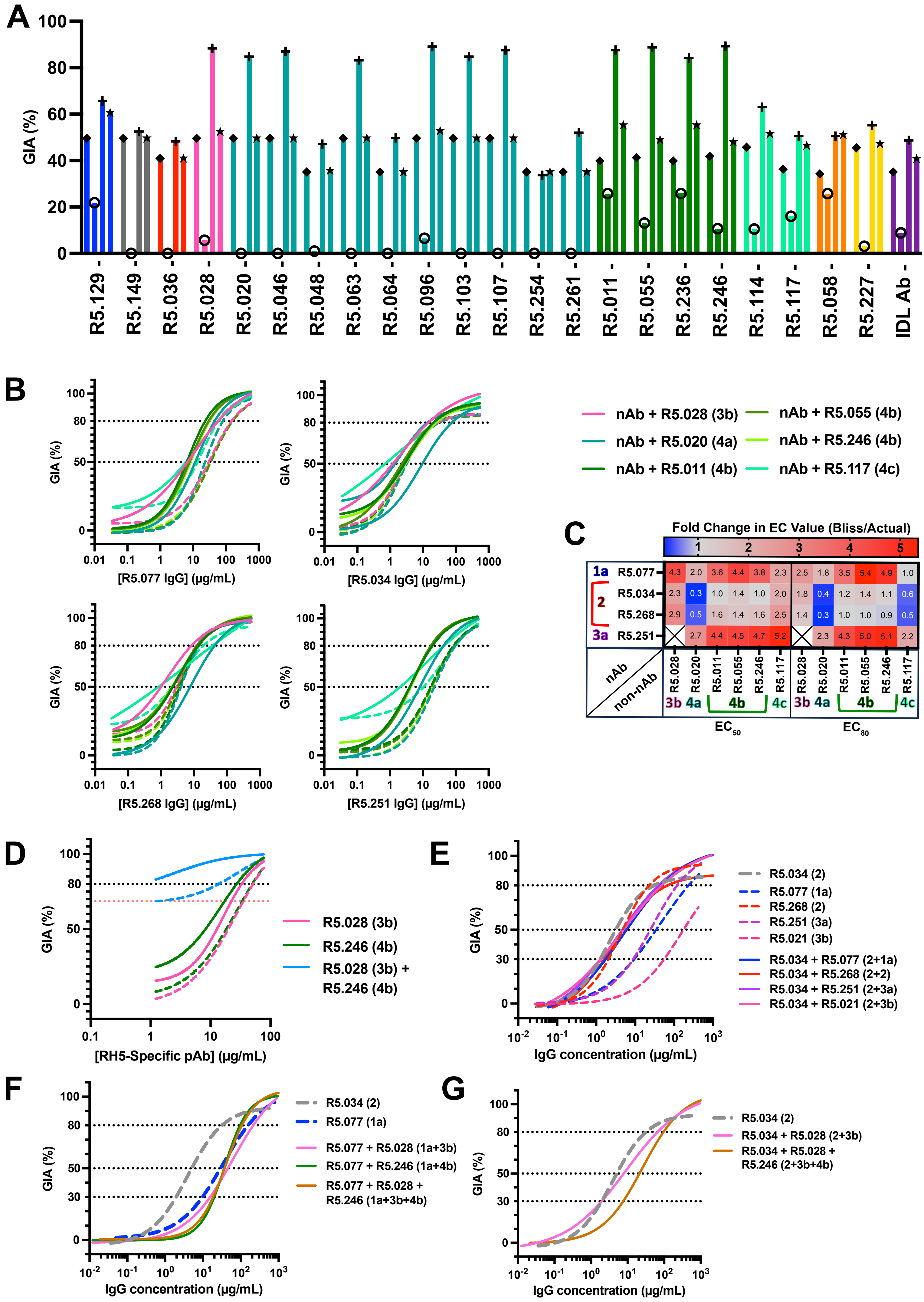
A
GIA (%)
0
20
40
60
80
100
R5.129
R5.149
R5.036
R5.028
R5.020
R5.046
R5.048
R5.063
R5.064
R5.096
R5.103
R5.107
R5.254
R5.261
R5.011
R5.055
R5.236
R5.246
R5.114
R5.117
R5.058
R5.227
IDL Ab
B
GIA (%)
[R5.077 IgG] (μg/mL)
[R5.034 IgG] (μg/mL)
[R5.268 IgG] (μg/mL)
[R5.251 IgG] (μg/mL)
0.01
0.1
1
10
100
1000
nAb + R5.028 (3b)
nAb + R5.020 (4a)
nAb + R5.011 (4b)
nAb + R5.055 (4b)
nAb + R5.246 (4b)
nAb + R5.117 (4c)
C
Fold Change in EC Value (Bliss/Actual)
1
2
3
4
5
1a R5.077 4.3 2.0 3.6 4.4 3.8 2.3 2.5 1.8 3.5 5.4 4.9 1.0
2 R5.034 2.3 0.3 1.0 1.4 1.0 2.0 1.8 0.4 1.2 1.4 1.1 0.6
R5.268 2.9 0.5 1.6 1.4 1.6 2.5 1.4 0.3 1.0 1.0 0.9 0.5
3a R5.251 2.7 4.4 4.5 4.7 5.2 2.3 4.3 5.0 5.1 2.2
nAb
non-nAb
R5.028 R5.020 R5.011 R5.055 R5.246 R5.117 R5.028 R5.020 R5.011 R5.055 R5.246 R5.117
3b 4a 4b 4c 3b 4a 4b 4c
$EC_{50}$
$EC_{80}$
D
GIA (%)
[RH5-Specific pAb] (μg/mL)
R5.028 (3b)
R5.246 (4b)
R5.028 (3b) + R5.246 (4b)
E
GIA (%)
IgG concentration (μg/mL)
R5.034 (2)
R5.077 (1a)
R5.268 (2)
R5.251 (3a)
R5.021 (3b)
R5.034 + R5.077 (2+1a)
R5.034 + R5.268 (2+2)
R5.034 + R5.251 (2+3a)
R5.034 + R5.021 (2+3b)
F
GIA (%)
IgG concentration (μg/mL)
R5.034 (2)
R5.077 (1a)
R5.077 + R5.028 (1a+3b)
R5.077 + R5.246 (1a+4b)
R5.077 + R5.028 + R5.246 (1a+3b+4b)
G
GIA (%)
IgG concentration (μg/mL)
R5.034 (2)
R5.034 + R5.028 (2+3b)
R5.034 + R5.028 + R5.246 (2+3b+4b)

### Figure S5: Functional assessment of intra-PfRH5 antibody interactions and combinations; related to Figure 5.

(**A**) Example GIA data for single concentration synergy screening of the community 1a nAb R5.077 in combination with non-nAbs. The x-axis displays the non-nAb used in combination and bars are colored according to the non-nAb's epitope community. Each bar is marked by a symbol. Diamond is for the GIA of R5.077 alone; hollow circle is for the non-nAb alone; plus is for the combination; and the star is for the Bliss additivity predicted GIA. R5.077 was tested at the $EC_{50}$ value of 15.6 µg/mL. Non-nAbs were tested at 0.3 mg/mL fixed final concentration. (**B**) Representative nAbs from communities 1a (R5.077), 2 (R5.034, R5.268) and 3a (R5.251) were tested in combination with a fixed concentration of representative modulatory non-nAbs from communities 3b (R5.028), 4a (R5.020), 4b (R5.011, R5.055 and R5.246) and 4c (R5.117). nAbs were tested in a 5-fold dilution series from a 0.5 mg/mL starting concentration. Non-nAbs were added at a final fixed concentration of 0.2 mg/mL to each point on the dilution curve. The data were log transformed and fitted to a four-parameter nonlinear regression model and EC values were interpolated. Horizontal dotted lines indicate 50 % and 80 % GIA, and predicted Bliss additivity GIA curves for each combination are shown as a dashed line. (**C**) Heat map of fold changes in $EC_{50}$ and $EC_{80}$ values for antibody pairs in **B**. The ratio of the Bliss additivity predicted EC value over the measured EC value was used to calculate fold change. A larger fold change indicates an improvement in the EC values, with values <1 being antagonistic (blue) and values >1 being synergistic (red) interactions, as indicated by the color scale. (**D**) GIA assay dilution curves of total IgG purified from the sera of RH5.1/$AS01_B$ vaccinees [1] run in a 2-fold dilution starting from ~14.4 mg/mL under various test conditions. PfRH5-specific IgG concentration within the purified total IgG was determined by quantitative ELISA [1] and used to plot the data on the x-axis. Data were also log transformed and a four-parameter non-linear regression was plotted. For each curve, a non-nAb (R5.028 from community 3b or R5.246 from community 4b), or a combination of both non-nAbs was added at a fixed concentration of 0.2 mg/mL each. Predicted Bliss additivity GIA curves for each combination are shown as a dashed line. The black dotted line indicates 80 % GIA. The red dotted line indicates the level of GIA measured alone for the fixed concentration combination of the two non-nAbs (R5.028+R5.246). (**E**) GIA titration curves of the highly potent R5.034 public clonotype mAb from community 2 tested in combination with other nAbs. The epitope community of each mAb clone is identified in parentheses. X-axis concentration indicates the total concentration of IgG in the test sample. Mixtures of antibodies (solid lines) were combined in equal ratios and compared to single mAb clones (dashed lines). The GIA curve for R5.034 is shown by a bold grey dashed line for ease of reference. Dotted lines indicate 30 %, 50 % or 80 % GIA. (**F**) Same experimental setup as **E** but testing R5.034 (best nAb in community 2) or (**G**) R5.077 (best nAb in community 1a) with the synergistic non-nAb clones R5.028 (community 3b) and/or R5.246 (community 4b).

Figure S6.

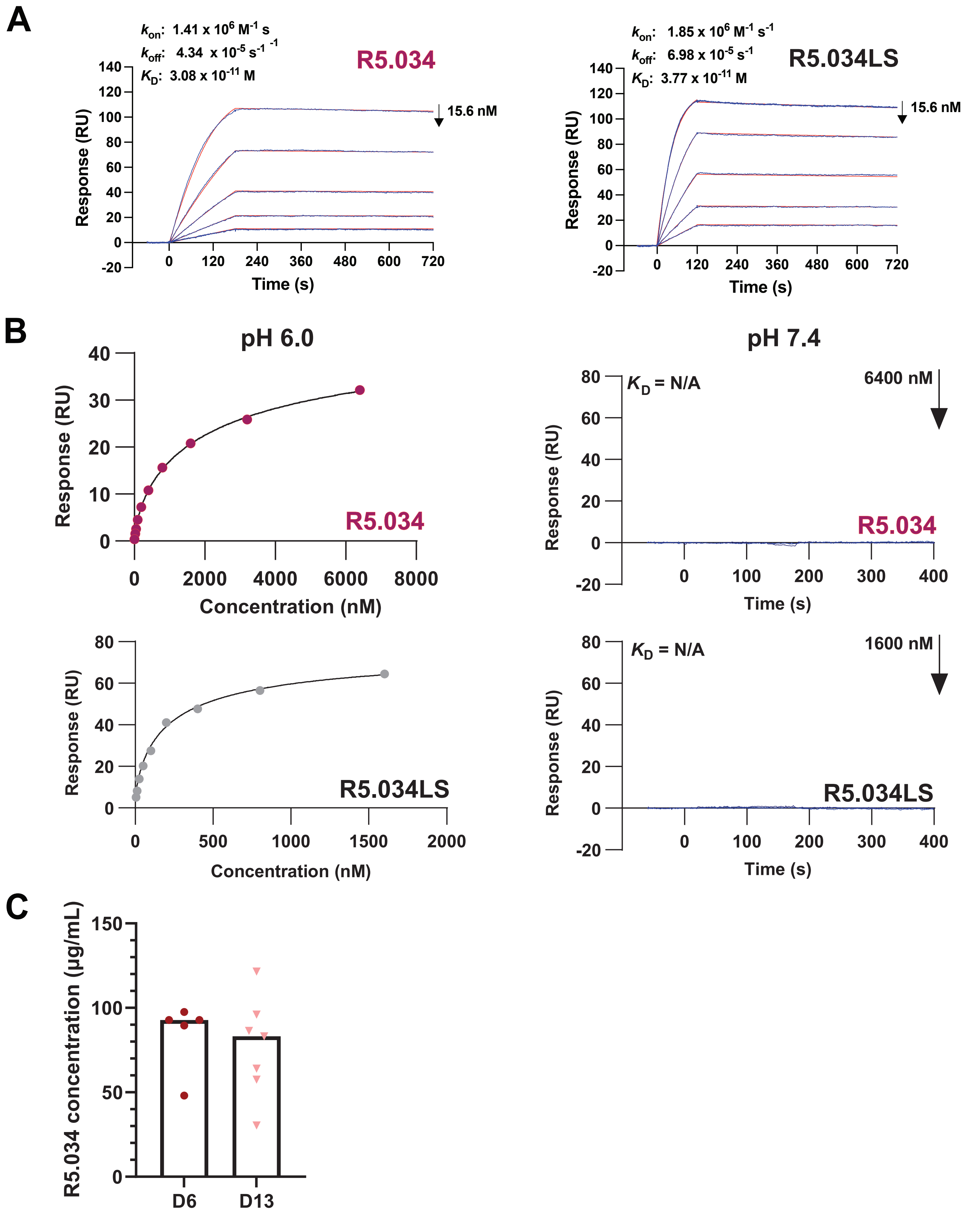

A
$k_{on}$: 1.41 x 10$^6$ M$^{-1}$ s$^{-1}$
$k_{off}$: 4.34 x 10$^{-5}$ s$^{-1}$
$K_D$: 3.08 x 10$^{-11}$ M
R5.034
15.6 nM
Response (RU)
Time (s)
$k_{on}$: 1.85 x 10$^6$ M$^{-1}$ s$^{-1}$
$k_{off}$: 6.98 x 10$^{-5}$ s$^{-1}$
$K_D$: 3.77 x 10$^{-11}$ M
R5.034LS
15.6 nM
Response (RU)
Time (s)
B
pH 6.0
Response (RU)
Concentration (nM)
R5.034
pH 7.4
$K_D$ = N/A
6400 nM
Response (RU)
R5.034
Time (s)
Response (RU)
R5.034LS
Concentration (nM)
$K_D$ = N/A
1600 nM
Response (RU)
R5.034LS
Time (s)
C
R5.034 concentration (μg/mL)
D6
D13

#### Figure S6: Binding affinities of R5.034 and R5.034LS, and mouse challenge ELISA; related to Figure 6.

(**A**) Association rate ($k_{on}$), dissociation rate ($k_{off}$) and affinity ($K_D$) values are shown as assessed by SPR. R5.034 or R5.034LS mAbs were immobilized and RH5.1 protein was injected in a 5-fold dilutions series from 15.6 nM concentration. Sensorgrams (blue) and fitted model (red) are shown with the data fit to the 1:1 Langmuir model. (**B**) Steady-state affinity, as assessed using SPR, of the R5.034 and R5.034LS mAb binding to human FcRn. Analysis at pH 6.0 (left hand panels, and sensorgrams shown in **Figure 6C**) or at pH 7.4 (right hand panels). Report points of the equilibrium binding levels at pH 6.0 of the 9-step 2-fold dilutions are fitted to the Equilibrium or Steady State binding model for $K_D$ determination. (**C**) Serum R5.034 antibody concentration in FRG huHep mice challenged with N54 strain *P. falciparum* was assessed by ELISA on days 6 (N=5) and 13 (N=7) post-challenge. The experiment was also terminated on day 13. Available data from individual mice are shown and the bar shows the median. ELISA data for the control group were all negative (data now shown).

### SUPPLEMENTAL TABLE

#### Table S1. Diffraction data collection and refinement statistics; related to Figures 4 and 6.

| | RH5ΔNL:R5.034 | RH5ΔNL:R5.251 |
|---|---|---|
| **PDB code** | 8QKS | 8QKR |
| **DATA COLLECTION** | | |
| **Wavelength (Å)** | 0.99987 | 0.9762 |
| **Space group** | $P2_1$ | P1 |
| **Cell dimensions** | | |
| a, b, c (Å) | 82.26, 376.79, 226.82, | 69.93, 92.02, 92.49, |
| α, β, γ (°) | 90.00, 90.06, 90.00 | 60.53, 72.13, 79.82 |
| **Resolution range** | 65.60 - 3.99 (3.99 - 3.99) | 59.11 - 3.29 (3.23 - 3.23) |
| **Total observation** | 779 885 (30 383) | 111 799 (5 486) |
| **Unique reflections** | 115 245 (5 241) | 30 110 (1 432) |
| ***R* pim (%)** | 0.176 (2.963) | 0.21 (1.17) |
| ***R* merge (%)** | 0.422 (6.530) | 0.35 (1.98) |
| ***R* meas (%)** | 0.458 (7.198) | 0.41 (2.30) |
| **$CC_{1/2}$** | 0.993 (0.220) | 0.955 (0.245) |
| **I / σ(I)** | 2.7 (0.1) | 3.1 (0.4) |
| **Completeness (%)** | 99.0 (90.0) | 98.6 (90.3) |
| **Multiplicity** | 6.8 (5.8) | 3.7 (3.8) |
| **Wilson B-factor** | 102.040 | 52.670 |
| **REFINEMENT** | | |
| **$R_{work}$** | 0.397 | 0.268 |
| **$R_{free}$** | 0.423 | 0.330 |
| **Average B, all atoms ($Å^2$)** | 93.0 | 76.0 |
| **Atoms, Protein** | 11747 | 32498 |
| **Atoms, Water** | 0 | 0 |
| **RMSD bonds** | 0.0146 | 0.0121 |
| **RMSD angles** | 1.65 | 2.0310 |
| **Ramachandran favoured (%)** | 76.5 | 91.7 |
| **Ramachandran allowed (%)** | 17.6 | 6.3 |
| **Ramachandran outliers (%)** | 5.9 | 1.9 |
| **Rotamer outliers** | 10.6 | 17.7 |
| **Clash score** | 19.6 | 9.4 |
| **MolProbity score** | 3.35 | 2.94 |

### SUPPLEMENTAL ITEMS

#### Data S1; related to Figures 1 and S1.

Supplemental data tables relating to **Figure 1** and **Figure S1**. (**A**) Data table of clones excluded from the epitope binning analysis with reasons for exclusion. (**B**) Normalized competition binning matrix for the epitope binning analysis. Squares in red indicate competition and squares in blue indicate sandwiching interactions. (**C**) Table summarizing the relationship between communities and supercommunities from the epitope binning analysis. (**D**) Table of GIA data values of all mAbs tested. Columns B-C relate to single concentration screening data. Columns E-S relate to data from GIA titration curves and interpolated values from the four-parameter fitted curves, including upper and lower 95% confidence intervals.

#### Data S2; related to Figure 2.

Full database of kinetic parameters relating to the kinetic data presented in **Figure 2**. Data represent the mean of N repeats as indicated in column AB.

#### Data S3; related to Figures 3 and S3.

Data relating to the sequence analysis performed in **Figure 3** and **Figure S3**. (**A**) Table of IMGT V-QUEST output parameters for mAb heavy and light chains. (**B**) Summary tallies of various gene and gene family usage and their combinations. (**C**) R Markdown file for the prediction of GIA from gene usage shown in **Figure S3F**.

#### Data S4; related to Figures 4 and 6.

Interfacing residues for the structural analysis performed in **Figure 4** and **Figure 6**. Table lists interfacing residues between PfRH5 (PDB ID: 4WAT numbering [6]) and the heavy or light chain of the indicated antibody. Interfacing residues were predicted by PDBePISA.
